# ER fusogens maintain membrane reservoir to ensure brain function

**DOI:** 10.1101/2025.04.12.648519

**Authors:** Hanchi Miao, Yuqi Wang, Yingying Huang, Haoyang Zhang, Lexin Zheng, Xin Zhou, Xiaoqun Wang, Qing Xie, Lei Xue, Junjie Hu

**Author notes:** These authors contributed equally (Hanchi Miao, Yuqi Wang). Correspondence (J.H.) and (L.X.).

## Abstract

How the morphological dynamics of the endoplasmic reticulum (ER) are linked to its functions is unclear. Atlastins (ATLs), a class of GTPases, mediate ER fusion, and human mutations in ATL1 cause hereditary spastic paraplegia (HSP). Here, we show that ATL2-knockout mice are embryonic lethal with compromised development, particularly of the cerebellum. ATL2 is highly expressed in neuroglia, though ATL1 is dominant in the brain. Lack of ATL2 disorganizes the positioning of Bergmann glia, which in turn interferes with granule cell migration. These cells have significant shrinkage in the intracellular membrane area, which is associated with decreased phosphatidylcholine and cholesterol synthesis. When tested in calyx-type synapses in ATL-deleted mice, a reduced membrane reservoir, represented by fewer presynaptic vesicles, leads to defective synaptic function and deafness. Collectively, these findings suggest that ER-shaping activity by ATL is essential for sustained lipid synthesis, and boosted lipid uptake is potentially beneficial for HSP patients.

## Introduction

Membrane fusion is a fundamentally important process involved in membrane trafficking and viral invasion (Jahn et al., 2003). Cellular organelles also undergo frequent fusion. Merging of mitochondria is vital for their fitness and genomic stability (Gao and Hu, 2021)(Veatch et al., 2009). Formation of the characteristic network of the endoplasmic reticulum (ER) via membrane fusion relies on a class of dynamin-like membrane-bound GTPases, namely atlastins (ATLs) (Hu et al., 2009). Being a fundamental cellular process, fusion of ER membranes is conserved throughout eukaryotic cells. Such activity is maintained in yeast by Sey1p and in plants by ROOT HAIR DEFECTIVE 3 (RHD3), both of which are functional analogs of ATL. In many species, such as *Drosophila melanogaster*, *Caenorhabditis elegans,* and *Danio rerio*, only one ATL is expressed. In rodents and higher mammals, three ATLs have been found: ATL1, ATL2, and ATL3.

Deletion of Sey1p along with an ER-localized SNARE, Ufe1p, causes synthetic lethality of *Saccharomyces cerevisiae* (Anwar et al., 2012). Lack of RHD3 leads to slowed growth of *Arabidopsis thaliana* with short and wavy root hairs, and additional loss of either RHD3-like protein renders lethality (Zhang et al., 2013). Deletion or depletion of ATL in model organisms that contain a single copy of ATL causes a variety of developmental defects (Orso et al., 2009)(Klemm et al., 2013)(Fassier et al., 2010). Importantly, mutations in ATL1 are frequently identified in patients with hereditary spastic paraplegia (HSP), a neurodegenerative disease characterized by progressive spasticity and weakness of the lower limbs due to a length-dependent abnormality of corticospinal axons (Salinas et al., 2008). Collectively, these observations underscore the physiological significance of ATL family proteins.

The ATLs contain an N-terminal GTPase (G domain) and a three-helix bundle (3HB), followed by two closely spaced transmembrane (TM) segments and a C-terminal tail (CT) (Bian et al., 2011). GTP-dependent dimerization of the G domain and hydrolysis-induced conformational movements of the 3HB lay the molecular foundation for membrane tethering and subsequent fusion (Bian et al., 2011). An amphipathic helix (APH) following the TM segment plays an auxiliary role in the fusion reaction (Faust et al., 2015)(Liu et al., 2012). Recent studies have revealed that the sequence after the APH also has a regulatory effect on ATL functions (Crosby et al., 2022)(Jang et al., 2023)(Krishna and Ford, 2023).

In mammals, ATL1 is predominantly localized to the central nervous system (CNS), whereas ATL2 and ATL3 are expressed mostly in peripheral tissues (Rismanchi et al., 2008). Consistently, almost all HSP-causing mutations are found in ATL1, with some exceptions in ATL3, but not in ATL2 (Salinas et al., 2008)(Behrendt et al., 2019)(Behrendt et al., 2021). Given the high levels of sequence similarity and equivalent GTPase and fusion activity, the three ATLs likely have redundant roles. However, ATL3 is a weaker fusogen than ATL1 and ATL2 (Hu et al., 2015). Whether an individual ATL or the ATL family as a whole is essential has not been tested in animal models.

A variety of ER-related cellular processes have been directly or indirectly linked to ATL (Lü et al., 2020). These processes include membrane trafficking, lipid metabolism, autophagy, microtubule dynamics, pathogen infections, calcium signaling, and even protein homeostasis. Because ATL mediates membrane tethering of the ER, deletion of ATL has been reported to compromise lateral membrane tension, which in turn slows cargo mobility and subsequent ER export (Niu et al., 2019). Similarly, ATL activity has been implicated in regulating the size of lipid droplets (LDs) (Klemm et al., 2013). These data indicate that ATL touches on a broad range of cellular events, from protein to lipid homeostasis. However, the physiological roles of ATLs are yet to be determined. Through a series of ATL knockout (KO) mice, we found that ATL2 plays an essential role in the maintenance of lipid synthesis that supports cerebellar development and synaptic function. In addition, combined ATL activity appears to be critical for brain cell viability.

## Results

### ATL2 is essential for cerebellar development

To investigate the physiological relevance of ER morphology, we generated whole-body KO mice of ATL1 and ATL2 (**Fig. S1A**). Although ATL1 KO mice are expected to exhibit symptoms of early onset HSP, they had no detectable developmental defects and showed a normal footprint pattern and walking track (**Fig. S1B**). To mimic the pathogenic conditions, in which mutated ATL1 may impose dominant-negative effects on remaining ATLs, we also generated ATL1 K80A knockin (KI) mice (**Fig. S1A**). K80 of ATL1 is critical for GTP interactions and its substitution with Ala results in significant reduction of the enzymatic activity (Yan et al., 2015). However, the KI mice behaved just like the wild-type (WT) mice in regard to brain development and walking track (**Fig. S1C**). In contrast, ATL2 KO mice were embryonic lethal (**Fig. S1A** and **Fig. 1A**), with developmental deficits in multiple organs, particularly a lack of brain tissue indicated by β-III-tubulin staining (**Fig. 1B** and **Fig. S1D,E**).

**Figure 1.**
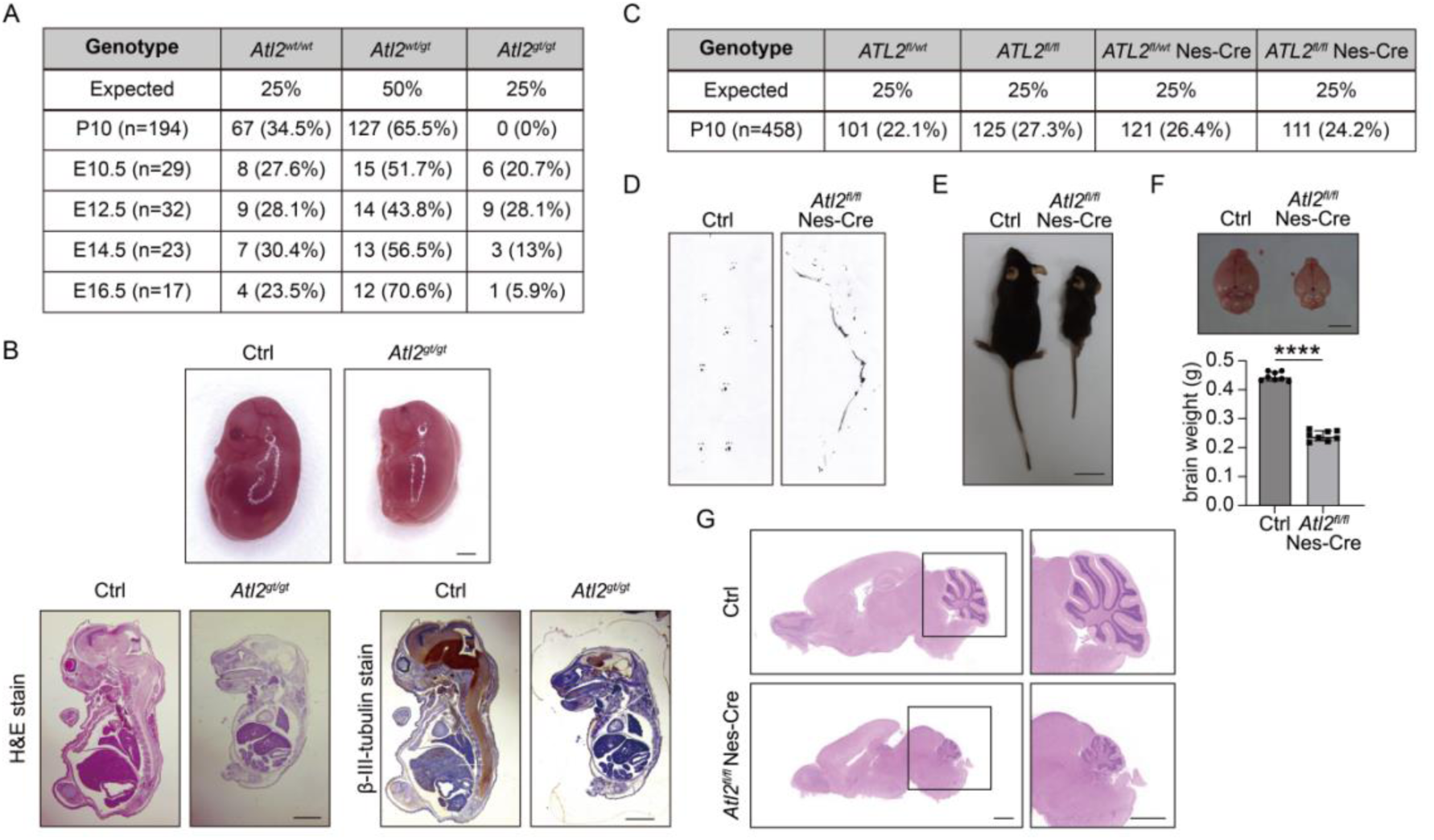
ATL2 is essential for cerebellar development. **A:** Distributions of offspring from intercrosses between *ATL2^gt/+^* mice. **B:** H&E staining and β-III-tubulin immunohistochemical staining of control and *ATL2^gt/gt^* embryos at E16.5 (scale bars, 2 mm). **C:** Distributions of offspring from intercrosses between *ATL2^fl/fl^* and *ATL2^fl/wt^* Nes-Cre mice. **D:** Footprint assay of control and *ATL2^fl/fl^* Nes-Cre mice. **E:** Image of control and *ATL2^fl/fl^*Nes-Cre mice (scale bar, 2 cm). **F:** Image and tissue weights of brains of control and *ATL2^fl/fl^* Nes-Cre mice (scale bar, 0.5 cm). **G:** H&E staining of the sagittal sections of brains from control and *ATL2^fl/fl^* Nes-Cre mice (scale bars, 1 mm). Data are represented as mean ± SD. The statistical significance of mean values was analyzed using an unpaired Student’s t test, \*\*\*\**p* < 0.0001.

ATL1 is the dominant ATL form in the CNS, but deletion caused no evident neurological problems. Therefore, we tested whether ATL2 plays an unexpected role in the CNS. Mice with conditional knockout (CKO) in the nervous system (ATL2^Nestin^-CKO) were successfully generated by crossing Nestin-Cre transgenic mice with ATL2^flox/flox^ mice (**Fig. S1A**). These mice were viable (**Fig. 1C**) but often suffered pup death due to shortage of movement-related food uptake (**Fig. S1F**). They had uncoordinated footprints (**Fig. 1D** and **movie S1**). The surviving mice had significantly reduced body size and weight compared to WT mice (**Fig. 1E** and **Fig. S1G**). As expected, the most severe developmental defects were in the brain (**Fig. 1F**). In particular, the cerebellum of these mice was significantly smaller (**Fig. 1G**), which explained their imbalanced movements. These results demonstrate that ATL2 is crucial for the growth of multiple organs and essential for cerebellar development in mice.

Next, we tested whether ATL2 is needed in adult mice. ATL2^flox/flox^ mice were crossed with transgenic mice expressing tamoxifen-inducible Cre recombinase, and tamoxifen was injected when the mice were 2 months old. Two months after the treatment, we noticed specific shrinkage in the cerebellum area (**Fig. S1H**) without a detectable loss in body weight (**Fig. S1I**). These results demonstrate that ATL2 also supports cerebellum health in adult mice, but its absence in adulthood is not life-threatening.

### ATL2 is highly expressed in the cerebellum and neuroglia

To test whether ATL2 dependencies are linked to its expression pattern, we systematically compared levels of ATLs in various tissues and regions. Consistent with previous reports (Rismanchi et al., 2008), ATL1 was expressed predominantly in brain tissue, whereas ATL2 and ATL3 were mostly found in peripheral tissues, with ATL2 being most abundant in muscle and ATL3 in lung (**Fig. 2A**). Because we observed a brain development phenotype in ATL2 KO mice, we focused on the brain. During embryonic development, ATL1 and ATL2 levels in the brain were gradually elevated, but ATL3 mostly remained constant (**Fig. 2B**), suggesting more important roles for ATL1 and ATL2 in the brain. Deletion of ATL2 in the nervous system did not trigger compensatory expression of ATL1 and ATL3 (**Fig. 2C**). Intriguingly, when brain regions were further analyzed, ATL1 was enriched in the cerebrum but ATL2 was highly expressed in the cerebellum (**Fig. 2D**), in line with the fact that ATL2 deletion caused severe defects in cerebellar development. These results were confirmed using human brain tissue lysates (**Fig. 2E**).

**Figure 2.**
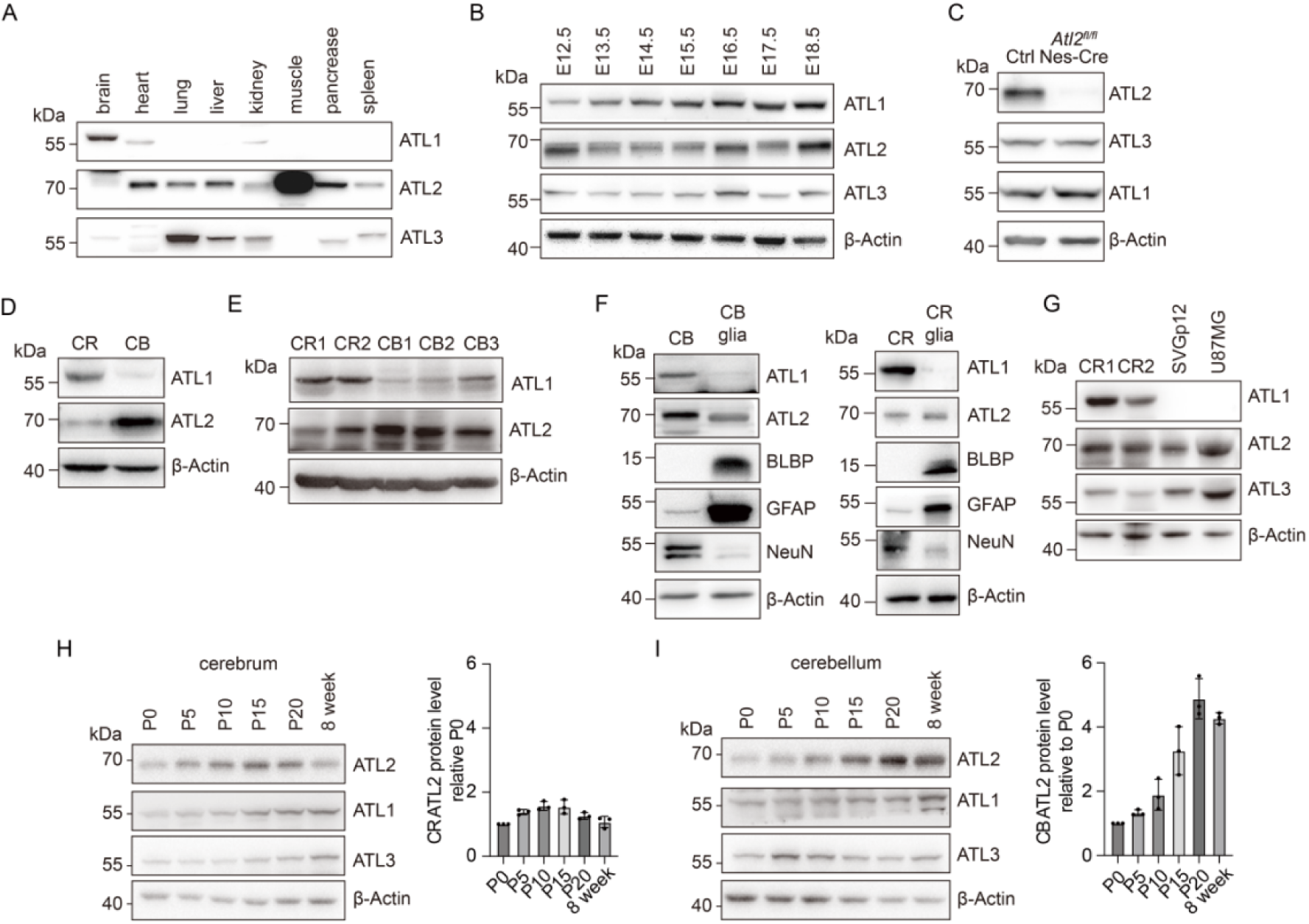
ATL2 is highly expressed in the cerebellum and neuroglia. **A-B:** Western blot analysis of the expression of three ATLs in different mouse tissues (A) and mouse brains from different embryonic stages (B). **C:** Western blot analysis of the expression of three ATLs in control and *ATL2^fl/fl^*Nes-Cre mice. **D-E:** Western blot analysis of ATL1 and ATL2 expression in mouse (D) and human (E) cerebrum and cerebellum. **F-G:** Western blot analysis of different ATLs in neuroglial cells from mouse (F) and human (G) brains. **H-I:** Western blot analysis of three ATLs in mouse cerebrum (H) and cerebellum (I) at different postnatal times. Data are represented as mean ± SD.

Both the cerebrum and cerebellum are composed of neurons and neuroglia. Next, we isolated these two brain regions and separated neuroglia for analysis. We found that ATL1 was largely missing from neuroglia, whereas ATL2 was abundant in these cells (**Fig. 2F**). We confirmed the fidelity of the isolation by blotting the neuron marker NeuN and glial markers BLBP and GFAP (**Fig. 2F**). When human brain and glial cell lines SVGp12 and U-87MG were analyzed, the same results were obtained (**Fig. 2G**). Finally, we monitored the expression pattern of ATL2 in the cerebrum and cerebellum during postnatal growth. Consistently, ATL2 levels slightly decreased in the cerebral tissue but evidently increased in the cerebellar region (**Fig. 2H,I**). In contrast, ATL1 gradually increased during cerebral development but exhibited only minor augmentation in the cerebellum (**Fig. 2H,I**). Similarly, ATL3 was increased in the cerebrum but had marginal fluctuations in the cerebellum (**Fig. 2H,I**). Our findings correlate with human and mouse brain transcriptome analyses (**Fig. S2A,B**) (Cardoso-Moreira et al., 2019). Taken together, these results suggest that ATL2 is indispensable for cerebellar development, possibly by supporting neuroglial function. ATL1 is likely important in neurons, but its role can be fulfilled by ATL2, or even ATL3.

### ATL activity is critical in all cerebellar cell types

To further dissect the specific role of ATL2 in cerebellar development, we focused on three major cell types in the cerebellum: Purkinje cells, granule cells, and glial cells (**Fig. 3A**). To this end, three CKO lines were generated by crossing ATL2^flox/flox^ mice with Pcp2-Cre (expressed in Purkinje cells), Atoh1-Cre (expressed in granule cells), or Gfap-Cre (expressed in neuroglia) transgenic mice. All three CKO mice were viable (**Fig. S3A-C**) and developed a regular-sized brain, but they developed a smaller cerebellum, though to differing extents (**Fig. 3A-C**). The body weights of the CKO mice were equivalent to WT, but their cerebellum organ indices were reduced (**Fig. S3D**). These results confirm that ATL2 contributes to all cell types in the cerebellum, with varied emphasis.

**Figure 3.**
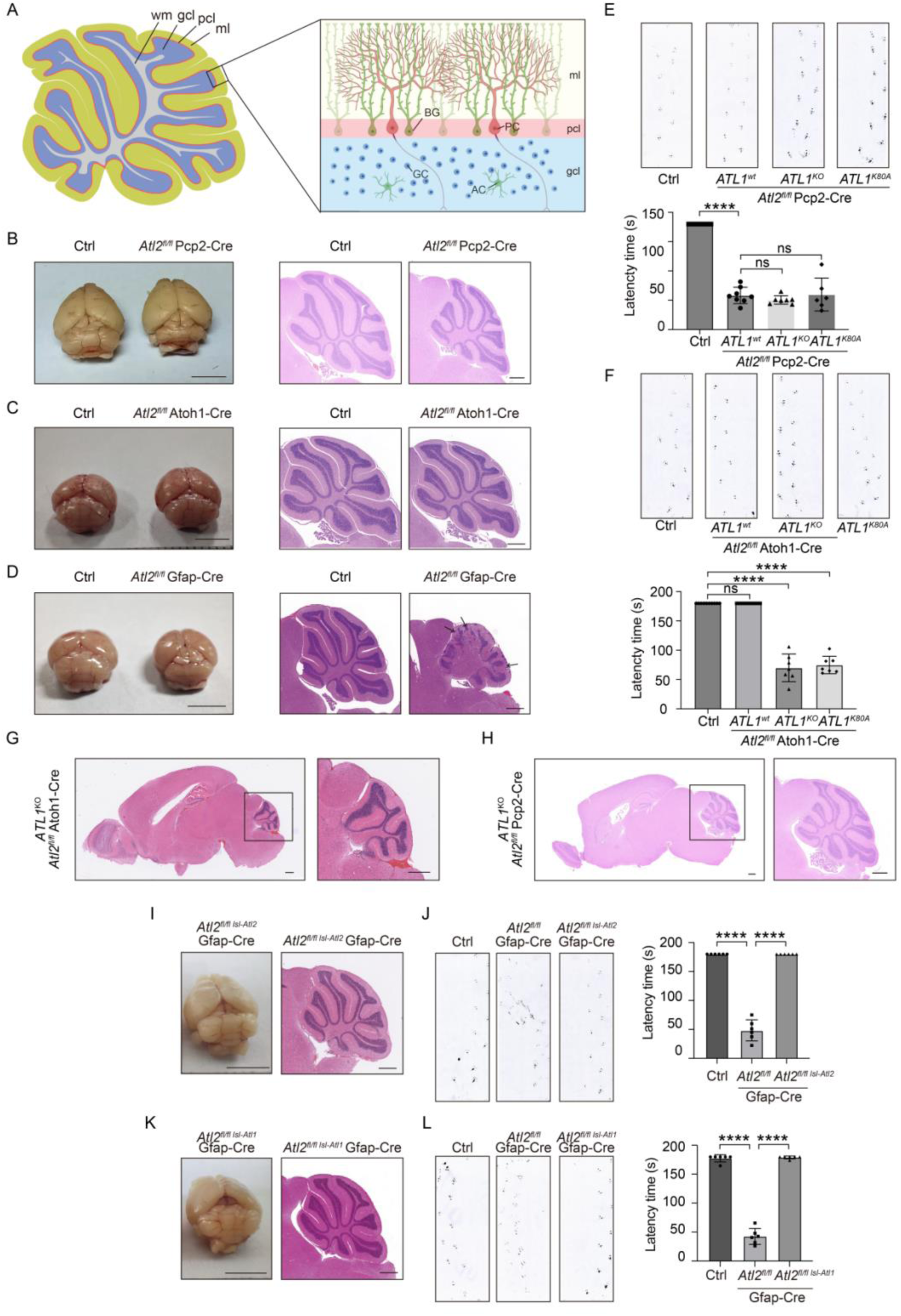
ATL activity is critical in all cerebellar cell types. **A:** Schematic of cerebellum structure. ml, molecular layer; pcl, Purkinje cell layer; gcl, granule cell layer; wm, white matter; BG, Bergmann glia; PC, Purkinje cell; AC, astrocyte; GC, granule cell. **B-D:** Brain image and cerebellum H&E staining of *ATL2^fl/fl^*Pcp2-Cre (B), *ATL2^fl/fl^* Atoh1-Cre (C), *ATL2^fl/fl^* Gfap-Cre (D), and their controls (scale bars, 0.5 cm in brain image, 0.5 mm in H&E staining). **E-F:** Footprint test (upper panel) and rotarod test (lower panel) of Purkinje neuron (E) and granule neuron (F) ATL1/2 double mutant mice. **G-H:** H&E staining of sagittal brain sections of granule neuron (G) and Purkinje neuron (H) ATL1/2 double knockout mice (scale bars, 0.5 mm). **I-L:** Brain images (left panel) and H&E staining of sagittal cerebellum sections (right panel) of *ATL2^fl/fl^* Gfap-Cre mice overexpressing ATL1 (I) or ATL2 (K) (scale bar, 0.5 cm in brain image, 0.5 mm in H&E staining). Footprint and rotarod test of *ATL2^fl/fl^*Gfap-Cre mice overexpressing ATL1 (J) or ATL2 (L). Data are represented as mean ± SD. The statistical significance of mean values was analyzed using an unpaired Student’s t test, \*\*\*\**p* < 0.0001, ns, not significant.

The ATL2^Pcp2^-CKO mice had a normal pattern in the cerebellum but it was only about half the size of the cerebellum found in the WT mice (**Fig. 3B**). When the organization and overall morphology of Purkinje cells were analyzed in the CKO mice, no detectable defects were found, but the number of Purkinje cells in the molecular layer was significantly reduced when stained by cell-specific marker calbindin (**Fig. S3F**). In addition, the area of the cerebellar cortex was evidently decreased in the CKO mice compared to WT mice, even though the cell density was not altered (**Fig. S3G**), suggesting a defect in cell proliferation. The ATL2^Pcp2^-CKO mice had normal footprints (**Fig. 3E**), but their latency time on a running wheel was reduced (**Fig. 3E**). These data suggest that ATL2 supports some cell proliferation mediated by Purkinje cells, but the outcome of ATL2 deletion in these cells is not as devastating as that of deletion of the whole nervous system.

The ATL2^Atoh1^-CKO mice also had cerebellar foliation similar to WT, with only a minor size reduction (**Fig. 3C** and **Fig. S3D**), suggesting less dependence on ATL2 in granule cells. As expected, the CKO mice exhibited normal footprints and running wheel performance (**Fig. 3F**). One plausible explanation is that ATL1 is sufficiently expressed in granule and Purkinje neurons, reducing the reliance of ATL2 in these cells. To test this possibility, we crossed ATL1 KO or K80A KI mice with ATL2^Pcp2^-CKO and ATL2^Atoh1^-CKO to generate ATL double-knockout (DKO) in either Purkinje or granule neurons. Both DKO lines were viable, and the cerebellar shrinkage and abnormality was exacerbated only in the Atoh1-related DKO mice (**Fig. 3G,H**). Consistently, the Atoh1-related DKO mice had defective running wheel capabilities (**Fig. 3E,F**). These results indicate that combined ATL activity is important, at least for granule cells, during cerebellar development.

Strikingly, the ATL2^Gfap^-CKO mice exhibited the most severe developmental and balancing defects among the three CKO lines, nearly reminiscent of ATL2^Nestin^-CKO. The organization of the cerebellum was severely damaged and the size significantly smaller (**Fig. 3D**). The organ index for the cerebellum was substantially decreased in these mice (**Fig. S3D**). In contrast, the organ indices of the cerebrum or whole brain were not affected (**Fig. S3E**), suggesting a much less dependable role of ATL2 in non-cerebellar neuroglia. As expected, the ATL2^Gfap^-CKO mice had imbalanced footprints and performed poorly in the running wheel tests (**Fig. 3J**). These results indicate that cerebellar defects caused by ATL2 deletion are most sensitive in cerebellar neuroglia, which correlates with the high and non-compensatory expression of ATL2 in these cells.

Next, we tested whether the role of ATL2 in neuroglia can be restored by any ATL. Canonical ATL2 (ATL2-1) has recently been reported to possess an auto-inhibitory effect, and splicing isoform ATL2-2, which is preferentially expressed in neuronal tissue, lacks the inhibition (Crosby et al., 2022). When the less active ATL2-1 was reintroduced into ATL2^Gfap^-CKO under the control of Gfap-Cre (**Fig. S3H**), cerebellar development was largely rescued (**Fig. 3I**), along with footprints and running wheel performance (**Fig. 3J**). Similarly, when ATL1 was expressed in Gfap-positive cells of ATL2^Gfap^-CKO, most defects were restored (**Fig. 3K,L**). When ATL2-1 or ATL1 was expressed in ATL2^Nestin^-CKO under the control of Nestin-Cre, they both efficiently rescued the phenotype seen with ATL2 deletion (**Fig. S3I,J**). Finally, when ATL2-1 or ATL1 was expressed in ATL2^gt/gt^ mice under the control of Cmv promoter, they both efficiently rescued the embryonic lethal phenotype (**Fig. S3K,L**). Taken together, these results suggest that the role of ATL2 in neuroglia can be fulfilled by other forms of ATL.

### ATL2 supports Bergmann glia-mediated granule cell migration

Next, we investigated specific cellular defects of cerebellar neuroglia that are associated with the developmental problem seen in the ATL2^Gfap^-CKO mice. When EAAT1, a glia-enriched transporter, was stained in the cerebella of WT or CKO mice, no apparent difference in signal strength was observed (**Fig. S4A**), suggesting that the overall amount of cerebellar glia is not changed between the two lines. However, when GFAP was stained to reveal intermediate filaments of cerebellar glia, significant disorganization was found in CKO mice. Specifically, the ladder-like pattern in the molecular layer was completely compromised in CKO cerebella (**Fig. 4A**). The ladder is composed of a type of neuroglia named Bergmann glia (BG), which serve as a scaffold for the radial migration of granule cells during laminar structure formation of the cerebellum (Xu et al., 2013). Notably, cells that failed to migrate and were trapped outside the cerebellar cortex were frequently spotted in both the ATL2^Nestin^-CKO and ATL2^Gfap^-CKO mice (**Fig. 1G** and **Fig. 3D**). Therefore, we performed an in vivo granule migration assay. As expected, granular migration was defective in the cerebellum of ATL2^Gfap^-CKO mice (**Fig. 4B** and **movie S2**). These results suggest that ATL2 deletion in BG causes its dysregulated positioning that leads to the most severe defects in cerebellar development.

**Figure 4.**
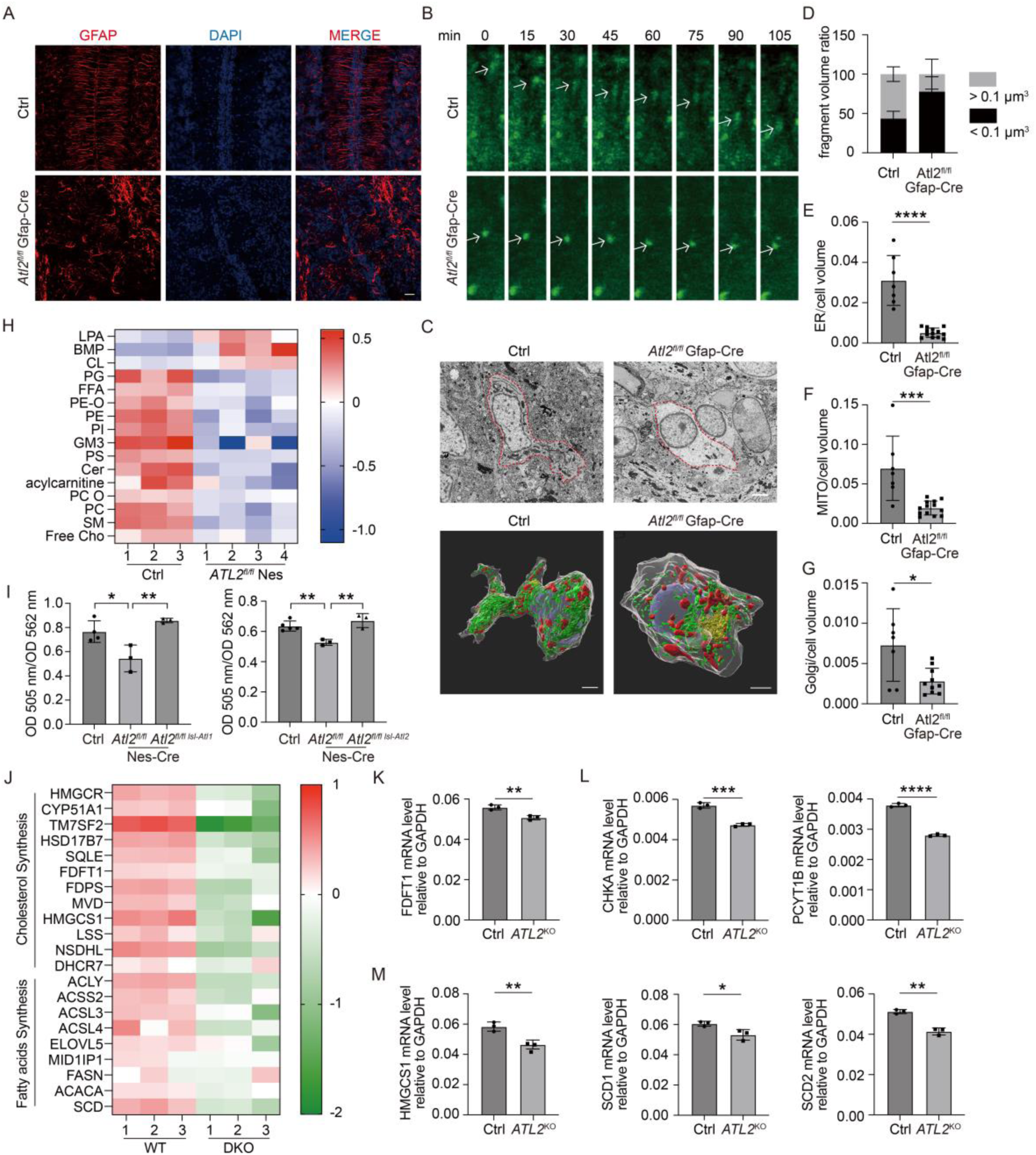
ATL2 deletion causes severe shrinkage of membrane reservoir. **A:** GFAP IF staining of cerebellum on postnatal day 7 (P7) in control and *ATL2^fl/fl^* Gfap-Cre mice (scale bar, 50 μm). **B:** Time-lapse imaging of granule neuron migration in the cerebellum cortex of P7 mice. **C:** FIB-SEM (upper panel) and 3D reconstruction (lower panel) of Bergmann glia from control and *ATL2^fl/fl^* Gfap-Cre mice (scale bar, 2 μm). **D:** Volume of endoplasmic reticulum fragment in control and *ATL2^fl/fl^* Gfap-Cre Bergmann glia. **E-G:** Ratio of total volume of endoplasmic reticulum (E), mitochondria (F) and Golgi (G) to cell volume in control and ATL2-deficient Bergmann glia. **H:** Lipidomics analysis of the cerebellum of control and *ATL2^fl/fl^* Nes-Cre mice. **I:** Relative cholesterol in the cerebellum of control, *ATL2^fl/fl^* Nes-Cre, and *ATL2^fl/fl^* Nes-Cre mice overexpressing ATL1 or ATL2. **J:** RNA-seq analysis of lipid synthesis-related genes in WT and ATL2/3 DKO COS7 cells. **K-M:** mRNA level of *FDFT1* (K), *CHKA, PCYT1B* (L), *HMGCS1, SCD1*, and *SCD2* (M) in primary neuron glia from control and *ATL2^fl/fl^* Nes-Cre mice. Data are represented as mean ± SD. The statistical significance of mean values was analyzed using an unpaired Student’s t test, \*\*\*\**p* < 0.0001, \*\*\**p* < 0.001, \*\**p* < 0.01, \**p* < 0.05.

### ATL2 deletion causes severe shrinkage of membrane reservoir

BG positioning requires precise guidance of cell adhesion and, as such, N-cadherin is enriched on the surface of these cells (Horn et al., 2018). Therefore, we tested whether ATL2 deletion hinders trafficking and subsequent surface presentation of N-cadherin. Protein levels of N-cadherin were not significantly altered in ATL2 CKO mice (**Fig. S4B**), and N-cadherin could be sufficiently detected on BG cells (**Fig. S4C**). Ectopic expression of N-cadherin in ATL2^Gfap^-CKO mice did not restore their developmental defects (**Fig. S4D**). Thus, ATL-regulated N-cadherin export likely does not determine the consequence of ATL2 deletion.

Because ATL is known to maintain the ER morphology by generating the characteristic tubular ER network (Zhao et al., 2016), we set out to analyze the ER structure and probe other potential structural changes in BG cells. Cerebellar sections were subjected to focused ion beam processing in combination with scanning electron microscopy (FIB-SEM). As expected, the ER in BG cells became less continuous when ATL2 was deleted (**Fig. 4C,D** and **movie S3-4**). The total volume of the ER was significantly reduced in the BG of ATL2^Gfap^-CKO mice (**Fig. 4E**). In addition, we found that the volumes of other organelles, including the Golgi and mitochondria, were also drastically reduced (**Fig. 4F,G**). We confirmed these findings using serial thin-sectioning electron microscopy (EM) with 3D reconstruction and obtained the same results with WT and ATL2-deleted BG cells (**Fig. S4E**). These data indicate that ATL2 deletion reduces the cellular membrane reservoir.

To test whether ATL deletion blocks lipid synthesis, we measured cellular lipid content using cells and tissues. As expected, ATL2/3 DKO COS-7 cells (an ATL-null cell, as no ATL1 expression is detected in COS-7 cells) (Niu et al., 2019) had significantly decreased levels of phosphatidylcholine (PC), the most abundant structural lipid in cells (**Fig. S5A**). In addition, cholesterol levels were much lower in these cells than in WT (**Fig. S5B**). Notably, these defects could not be recapitulated by transient knockdown (KD) of ATL2/3 (**Fig. S5C**), suggesting that the shortage of lipids is likely a consequence of accumulated and irreversible long-term effects.

When we isolated cerebella from WT or ATL2^Nestin^-CKO mice and subjected the tissue to lipidomics, similar results were obtained; a large subset of major lipids was down-regulated, including PC, phosphatidylethanolamine (PE), phosphatidylserine (PS), phosphatidylglycerol (PG), phosphatidylinositol (PI), and ceramide (Cer) (**Fig. 4H**). The cholesterol level was also reduced in the cerebella of these mice, but the defects were restored by expression of either ATL1 or ATL2 (**Fig. 4I**).

Reduced lipid levels can be caused by downregulation of lipid synthesis enzymes. We performed RNA-Seq analysis by comparing the WT and DKO cells. In general, many enzymes involved in cholesterol or fatty acid synthesis had decreased transcript levels (**Fig. 4J**). Based on the lipid analysis in these cells, key enzymes for PC and cholesterol synthesis were tested individually. As expected, FDFT1 (also known as squalene synthase) was significantly reduced at the protein and transcript levels in DKO cells (**Fig. S5D**). The same changes were observed in neuroglia isolated from ATL2^Nestin^-CKO mice (**Fig. 4K**), but not with transient KD of ATLs (**Fig. S5E**). We also tested key enzymes in de novo PC synthesis and found that choline kinase (CHKA) and phosphate cytidylyltransferase 1B (PCYT1B) were reproducibly down-regulated in DKO cells (**Fig. S5F**). Once again, these changes did not occur when ATLs were only transiently depleted (**Fig. S5G**), but they were observed in ATL2-deleted primary neuroglia (**Fig. 4L**). These results confirm that the lipid deficiency seen in ATL deletion is probably a long-term deficit and correlates with decreased expression of lipid-producing enzymes.

Next, we traced the cause of the reduced lipid synthesis in the prolonged absence of ATL. Lipid synthesis is known to be controlled by sterol regulatory-element binding proteins (SREBPs), with SREBP-1 being a master regulator for fatty acid (FA) and triacylglycerol synthesis and SREBP-2 specific to cholesterol synthesis (Amemiya-Kudo et al., 2002). Activation of SREBP requires ER-to-Golgi trafficking of the protein and its subsequent cleavage, releasing a soluble form for transcriptional regulation. The process is accompanied by SREBP cleavage-activating protein (SCAP). Thus, sterol-induced translocation of SCAP from the ER to Golgi is commonly used to indirectly measure the capacity of SREBP activation. As expected, SCAP efficiently re-localized to the Golgi upon sterol depletion in WT COS-7 cells, but not DKO cells (**Fig. S5H**). The levels of subsequent SREBP activation were confirmed by monitoring the transcriptional levels of FA synthase (FASN) (**Fig. S5J**). Similarly, compromised SREBP activation were seen in neuroglia isolated from ATL2^Nestin^-CKO mice (**Fig. 4M**). Consistently, transient depletion of ATLs was not sufficient to trigger these severe defects in cells (**Fig. S5I**). Because we previously found that lack of ATL delays ER protein export (Niu et al., 2019), we reasoned that the lipid shortage seen with ATL deletion is most likely caused by sustained trafficking inhibition of SCAP and the subsequent prevention of SREBP activation.

We tested whether LDs, the major form of storage of intracellular lipids, are also affected by ATL deletion. ATL may directly mediate fusion of LDs (Klemm et al., 2013). We confirmed that, in DKO cells, LDs became smaller but greater in number (**Fig. S5K**). Similarly, LDs in cerebellum-derived neuroglia were smaller in size and increased in number when ATL2 was deleted in these cells (**Fig. S5M**). However, when ATL was transiently depleted, the same results were obtained (**Fig. S5L**). These results indicate that ATL plays a role in LD homeostasis, but whether the mechanism lies in LD fusion or lipid supplies is not clear. In summary, sustained ATL activity ensures sufficient lipid synthesis.

### ATL regulates vesicle recycling in the central nervous system

Synaptic transmission relies on continuous vesicle exocytosis and endocytosis, processes essential for neuronal communication (Schweizer and Ryan, 2006). We reasoned that membrane reservoir shortage could be sensitive for synaptic transmission, as synaptic vesicle pool maintenance relies heavily on lipid supply. To address this, we examined the role of ATL in presynaptic function by analyzing vesicle exo-endocytosis at the calyx of Held, a giant glutamatergic synapse in the auditory brainstem, using patch-clamp recordings on postnatal days 7-10 (p7-p10) in WT and ATL2^Nestin^-CKO mice. Depolarization for 20 ms (–80 to +10 mV, depol_20ms_) induced calcium influx and exocytosis (measured as capacitance jump, ΔCm). Depol_20ms_ can deplete the readily releasable pool (RRP) and trigger clathrin-dependent slow endocytosis (Liu et al., 2019)(Wu et al., 2020). In ATL2 CKO mice, both calcium influx and exocytosis were significantly reduced compared to WT (**Fig. 5A-C**). The subsequent endocytosis, reflected by an exponential decay in the capacitance trace, was also impaired, with slower endocytosis rates (Rate_endo_; **Fig. 5D**) and higher residual capacitance 15 s after stimulation (ΔCm_15s_%; **Fig. 5E**). These results indicate that ATL2 deletion disrupts calcium influx, exocytosis, and clathrin-dependent endocytosis.

**Figure 5.**
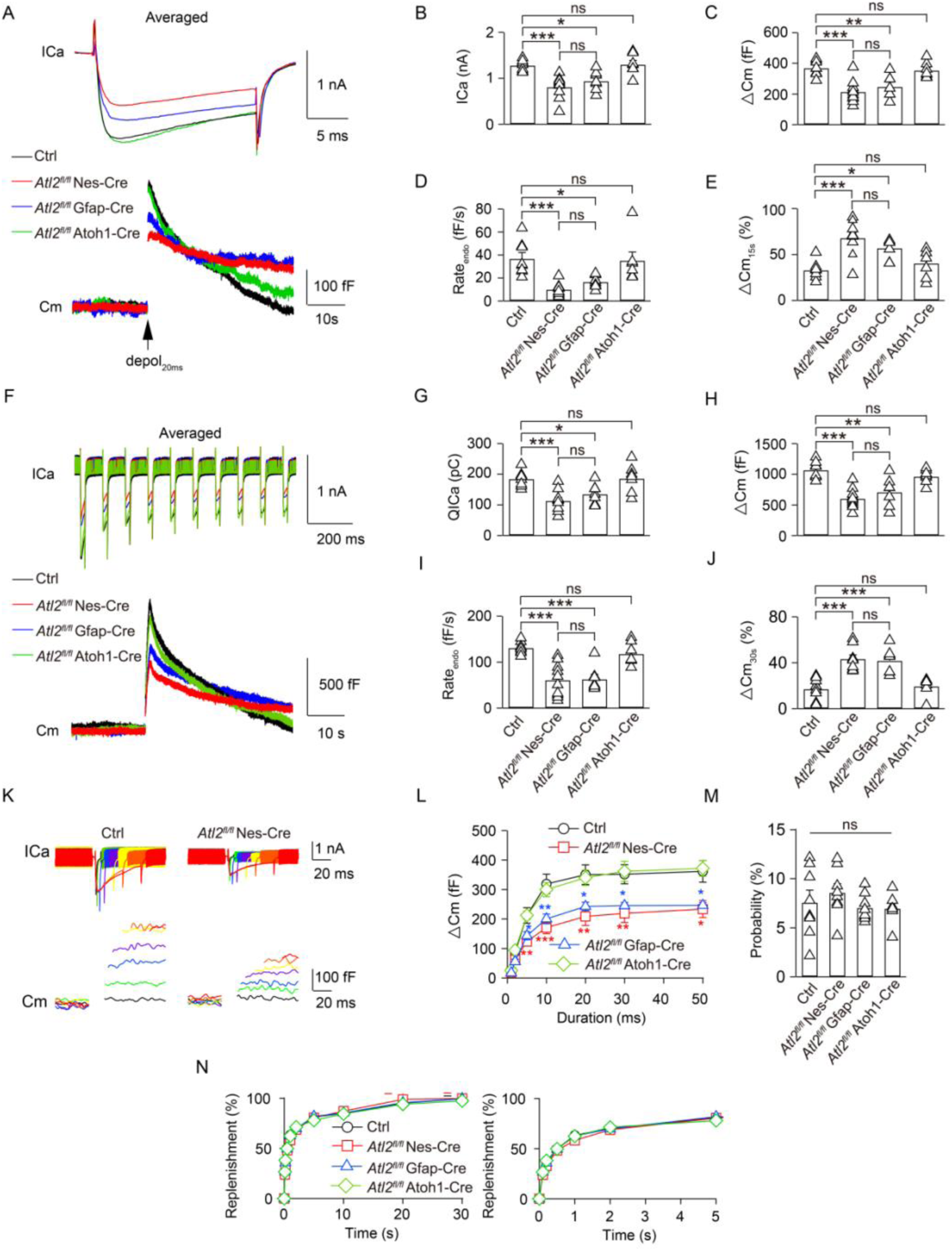
ATL2 deletion inhibits vesicle exo-endocytosis. **A**: Averaged presynaptic ICa (top) and Cm (bottom) induced by depol_20ms_ (arrow) in control (black), *ATL2^fl/fl^* Nes-Cre (red), *ATL2^fl/fl^* Gfap-Cre (blue), and *ATL2^fl/fl^* Atoh1-Cre (green) mice. **B-E**: Statistics for presynaptic ICa (B), ΔCm (C), Rate_endo_ (D), and ΔCm_15s_ (E) induced by depol_20ms_ from control (black), *ATL2^fl/fl^* Nes-Cre (red), *ATL2^fl/fl^* Gfap-Cre (blue), and *ATL2^fl/fl^*Atoh1-Cre (green) mice (control, n = 8; *ATL2^fl/fl^* Nes-Cre, n = 10; *ATL2^fl/fl^* Gfap-Cre, n = 7; *ATL2^fl/fl^* Atoh1-Cre, n = 7). **F**: Averaged presynaptic ICa (top) and Cm (bottom) induced by depol_20msx10_ (arrow) in the control (black), *ATL2^fl/fl^*Nes-Cre (red), *ATL2^fl/fl^* Gfap-Cre (blue), and *ATL2^fl/fl^*Atoh1-Cre (green) mice. **G-J**: Statistics for presynaptic QICa (G), ΔCm (H), Rate_endo_ (I), and ΔCm_30s_ (J) induced by depol_20msx10_ from control (black), *ATL2^fl/fl^* Nes-Cre, *ATL2^fl/fl^* Gfap-Cre, and *ATL2^fl/fl^* Atoh1-Cre (green) mice (control, n = 8; *ATL2^fl/fl^* Nes-Cre, n = 10; *ATL2^fl/fl^* Gfap-Cre, n = 7; *ATL2^fl/fl^* Atoh1-Cre, n = 7). **K**: Sampled ICa (top) and Cm (bottom) induced by 1 (black), 2 (green), 5 (blue), 10 (purple), 20 (yellow), 30 (orange), and 50 ms (red) depolarization pluses from –80 to +10 mV in the control and *ATL2^fl/fl^*Nes-Cre mice. **L**: The relationship between ΔCm and the duration of depolarization pulses in the control (n = 8 for each data point; black), *ATL2^fl/fl^*Nes-Cre (n = 8 for each data point; red), *ATL2^fl/fl^* Gfap-Cre (n = 8 for each data point; blue), and *ATL2^fl/fl^* Atoh1-Cre (n = 6 for each data point; green) mice. **M**: Statistics for the release probability measured by the percentage of RRP release induced by a 1-ms depolarization pulse from –80 to +10 mV in the control (black), *ATL2^fl/fl^* Nes-Cre (red), *ATL2^fl/fl^*Gfap-Cre (blue), and *ATL2^fl/fl^* Atoh1-Cre (green) mice using one-way ANOVA with Dunnett’s *post hoc* test. ns, not significant. **N**: Cm induced by a 20-ms depolarization applied at various intervals after the conditional stimulus (depol_20ms_) in control (black), *ATL2^fl/fl^* Nes-Cre (red), *ATL2^fl/fl^* Gfap-Cre (blue), and *ATL2^fl/fl^* Atoh1-Cre (green) mice. Left and right panels show the same data at different scales. Data are represented as mean ± SEM. The statistical significance of mean values was analyzed using a one-way ANOVA with Dunnett’s *post hoc* test, \*\*\**p* < 0.001, \*\**p* < 0.01, \**p* < 0.05, ns, not significant.

To evaluate clathrin-independent endocytosis during intense stimulation, we applied depol_20msx10_ (10 pulses of depol_20ms_ at 10 Hz). In WT mice, this protocol induced a calcium influx (QICa) of 186 ± 16 pC, a total ΔCm of 979 ± 27 fF, and a Rate_endo_ of 129 ± 8 fF/s within 2 s after stimulation (**Fig. 5F-J**). In ATL2 CKO mice, the calcium influx, exocytosis, and endocytosis rates were all significantly reduced (**Fig. 5F-J**), confirming the role of ATL2 in both clathrin-dependent and clathrin-independent endocytosis. We further quantified the RRP size using variable pulse lengths (1, 2, 5, 10, 20, 30, and 50 ms). In ATL2 CKO mice, ΔCm was significantly smaller for stimuli between 5 and 50 ms, indicating a reduced RRP size while did not affect the release probability as the cause of impaired exocytosis (**Fig. 5K-M**). However, RRP replenishment at various intervals (Δt = 0.05–30 s) was unaffected (**Fig. 5N**). The changes in RRP size are in line with the findings that ATL2 deletion can cause shortage of the membrane reservoir.

Next, we measured whether ATL2 CKO mice possess a reduced RRP pool. Brain sections that contain the calyx of Held were isolated and subjected to EM analysis. As expected, the WT neuron contained large amounts of vesicles in the presynaptic regions, particularly the active zone (**Fig. 6A**). In contrast, ATL2^Nestin^-CKO and ATL2^Gfap^-CKO mice, but not ATL2^Atoh1^-CKO mice, had significantly fewer vesicles in the equivalent regions (**Fig. 6B-D**). These data demonstrate that the shrinkage of the membrane reservoir seen with ATL2 deletion mostly manifests as a reduced RRP pool in neurons.

**Figure 6.**
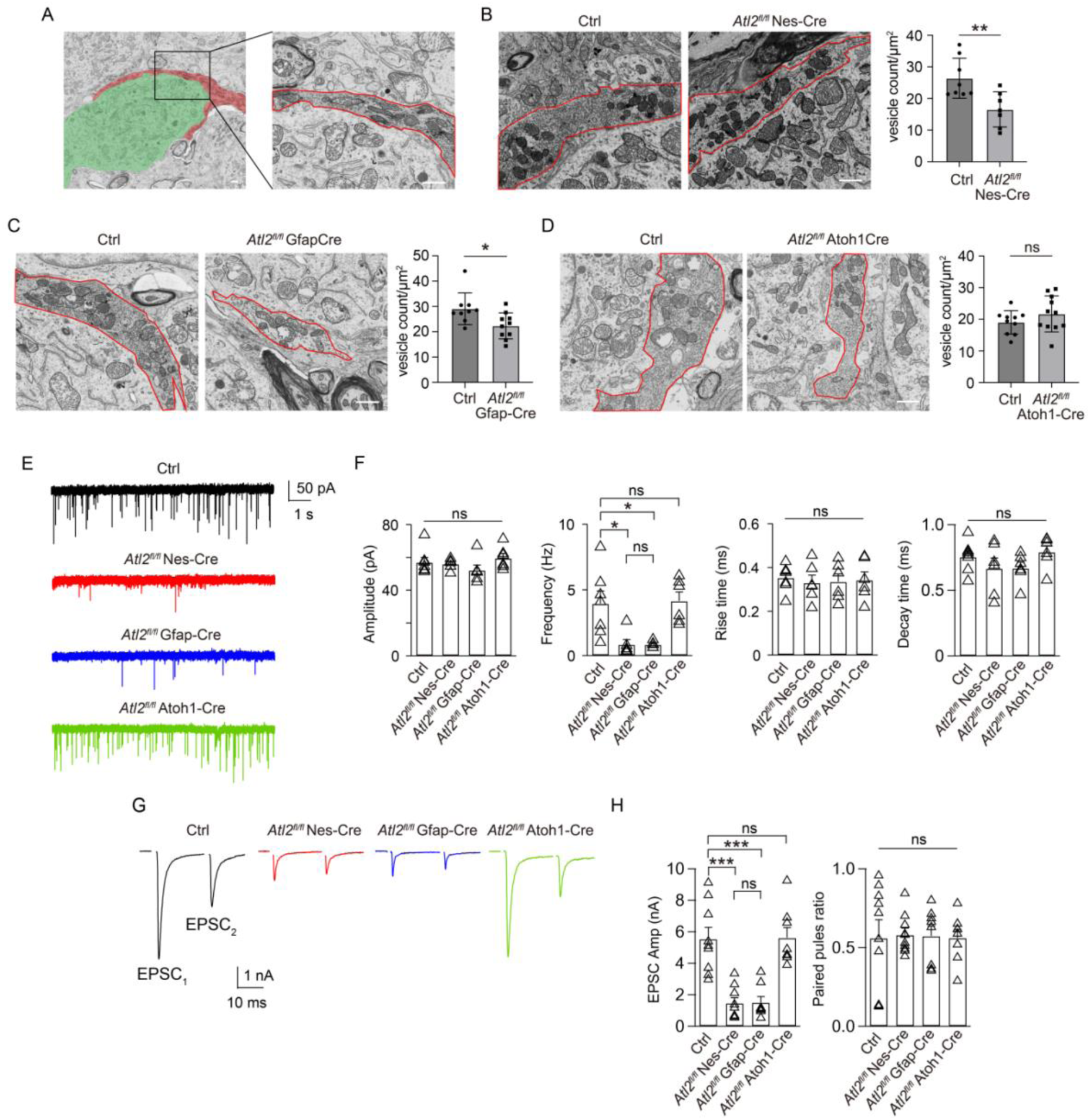
ATL2 deletion compromises synaptic transmission. **A:** EM image of red presynaptic calyx and green postsynaptic MNTB principal neuron (left). High-resolution EM images of the active zone (red box) within the calyx of Held (right) (scale bar, 1 μm). **B-D**: Synapse vesicle density of the calyx of Held from *ATL2^fl/fl^* Nes-Cre (B), *ATL2^fl/fl^* Gfap-Cre (C), and *ATL2^fl/fl^* Atoh1-Cre (D) mice and their controls. The red boxes indicate presynaptic active zones (scale bar, 1 μm). **E**: Representative traces of mEPSC recorded in control (black), *ATL2^fl/fl^* Nes-Cre (red), *ATL2^fl/fl^* Gfap-Cre (blue), and *ATL2^fl/fl^*Atoh1-Cre (green) mice. **F**: Statistics for the mEPSC amplitude, frequency, decay time, and rise time for all groups (control, n = 6; *ATL2^fl/fl^* Nes-Cre, n = 8; *ATL2^fl/fl^* Gfap-Cre, n = 6; *ATL2^fl/fl^* Atoh1-Cre, n = 8). **G**: Representative EPSC traces recorded in control (black), *ATL2^fl/fl^* Nes-Cre (red), *ATL2^fl/fl^* Gfap-Cre (blue), and *ATL2^fl/fl^* Atoh1-Cre (green) mice. **H**: Statistics for the EPSC amplitude and paired-pulse ratio (PPR) for all groups (control, n = 6; *ATL2^fl/fl^* Nes-Cre, n = 8; *ATL2^fl/fl^*Gfap-Cre, n = 6; *ATL2^fl/fl^* Atoh1-Cre, n = 8). Data in **B-D** are represented as mean ± SD. The statistical significance of mean values was analyzed using an unpaired Student’s t test, \*\**p* < 0.01, \**p* < 0.05, ns, not significant. Data in **F** and **G** are represented as mean ± SEM. The statistical significance of mean values was analyzed using one-way ANOVA with Dunnett’s *post hoc* test, \*\*\**p* < 0.001, \**p* < 0.05, ns, not significant.

We analyzed whether ATL2 is directly needed in neurons or indirectly involved in neuroglia. To test this, we confirmed that Atoh1-Cre is expressed in neurons at the calyx of Held (**Fig. S6A**). Neurotransmission was then assessed using ATL2^Atoh1^-CKO and ATL2^Gfap^-CKO mice. We found that ATL2^Gfap^-CKO mice phenocopied the defects seen in ATL2^Nestin^-CKO mice, whereas ATL2^Atoh1^-CKO mice behaved similarly as WT mice (**Fig. 5A-L**). These results confirm that ATL2 plays a more significant role in neuroglia than in neurons, influencing synaptic transmission and vesicle dynamics.

Because ATL1 is highly expressed in neurons, we tested whether ATL1 KO mice had similar problems in synaptic transmission. As expected, and in contrast to ATL2^Atoh1^-CKO mice, ATL1 KO mice recapitulated defects seen in ATL2^Nestin^-CKO and ATL2^Gfap^-CKO mice (**Fig. S6B-I**). Collectively, these findings highlight ATLs as critical regulators of vesicle recycling, affecting both clathrin-dependent and clathrin-independent endocytosis.

### ATL2 is essential for excitatory neurotransmission

To determine the role of ATL2 in excitatory neurotransmission, we recorded and compared miniature excitatory postsynaptic currents (mEPSCs) between WT and CKO mice. Although mEPSC amplitude, rise time, and decay time were similar across all groups, mEPSC frequency was significantly reduced in ATL2^Nestin^-CKO and ATL2^Gfap^-CKO mice compared to WT mice but remained unchanged in ATL2atoh1-CKO mice (**Fig. 6E,F**).

We further assessed EPSCs and paired-pulse ratios (PPRs) in WT and ATL2 CKO mice. EPSC amplitudes in ATL2^Atoh1^-CKO mice were comparable to those in WT mice but were significantly reduced in ATL2^Nestin^-CKO and ATL2^Gfap^-CKO mice (**Fig. 6G,H**). Despite the reduction in EPSC amplitude, PPRs did not differ significantly across groups, indicating that ATL2 deficiency decreases the availability of presynaptic vesicles without affecting vesicle release probability (**Fig. 6G,H**).

### ATL2 is essential for auditory function

To investigate the impact of ATL2 deficiency on auditory processing, we measured auditory brainstem responses (ABRs) in WT and ATL2 CKO mice (p20-p25) (Kim and Holt, 2013)(Jang et al., 2019). ABRs, which reflect synchronized neuronal activity in the auditory pathway, consist of five distinct peaks (waves I–V) in response to tonal stimuli (Jang et al., 2019). Representative waveforms at 16 kHz and 90 dB revealed rapid waveform flattening in ATL2^Nestin^-CKO and ATL2^Gfap^-CKO mice, indicative of elevated hearing thresholds compared to WT mice (**Fig. 7A**). Furthermore, the wave I amplitude was significantly reduced, and its latency was notably prolonged in the ATL2 CKO mice (**Fig. 7B**).

**Figure 7.**
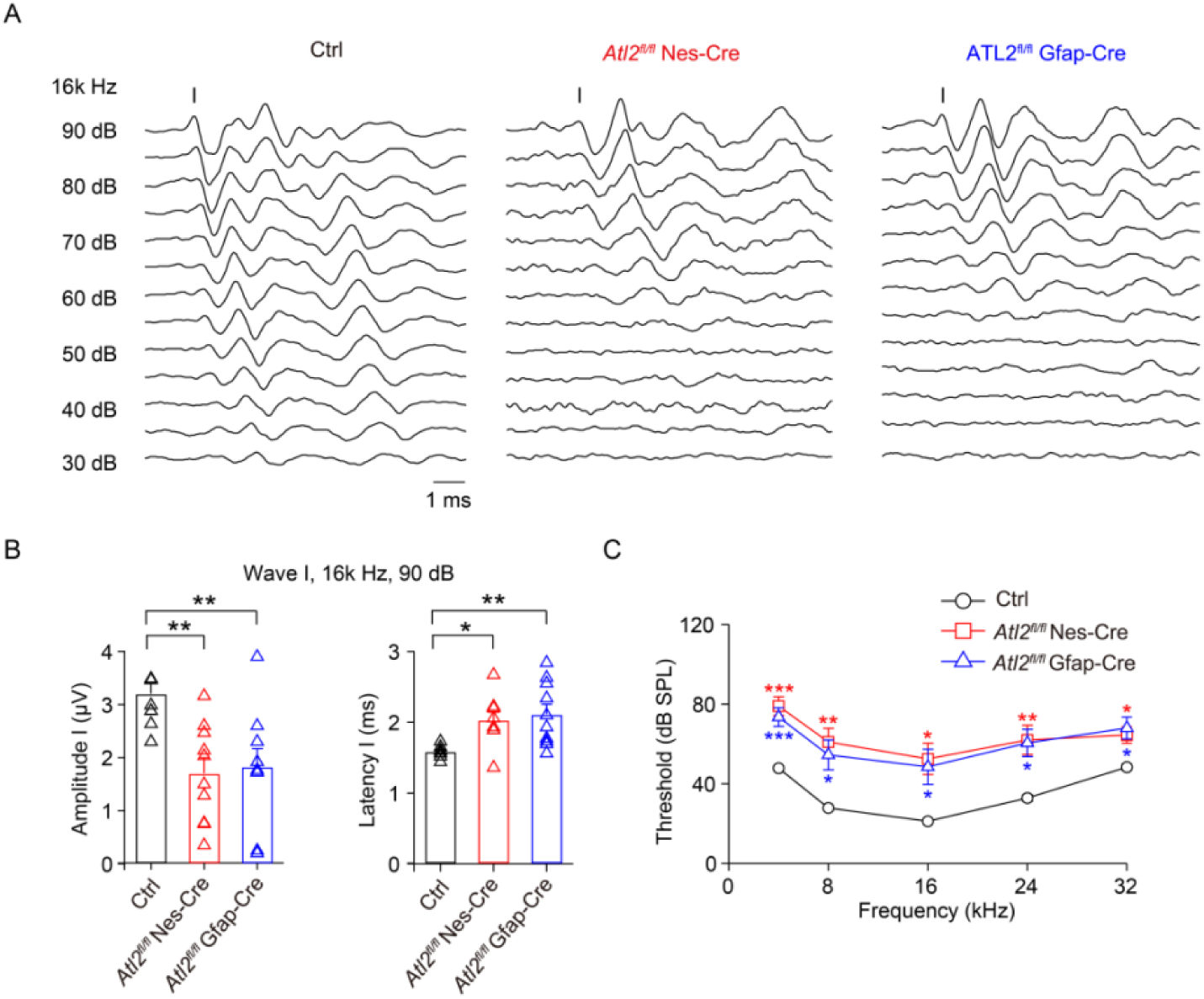
ATL2 deletion impairs auditory function. **A**: Representative ABR traces recorded using tone stimuli in control, *ATL2^fl/fl^*Nes-Cre, and *ATL2^fl/fl^* Gfap-Cre mice. Roman numerals identify wave I in control, *ATL2^fl/fl^* Nes-Cre, and *ATL2^fl/fl^*Gfap-Cre mice. **B**: Statistics for the amplitude and latency of wave I at 16 kHz and 90 dB in control, *ATL2^fl/fl^* Nes-Cre, and *ATL2^fl/fl^*Gfap-Cre mice. **C**: ABR thresholds at frequencies of 4, 8, 12, 24, and 32 kHz were recorded using tone stimuli in control, *ATL2^fl/fl^* Nes-Cre, and *ATL2^fl/fl^* Gfap-Cre mice. Data are represented as mean ± SEM. The statistical significance of mean values was analyzed using one-way ANOVA with Dunnett’s *post hoc* test, \*\*\**p* < 0.001, \*\**p* < 0.05, \**p* < 0.05.

We also assessed the frequency dependence of ABR thresholds using tone stimuli across a range of frequencies (4, 8, 12, 24, and 32 kHz) at varying intensities (90-30 dB). ATL2^Nestin^-CKO and ATL2^Gfap^-CKO mice displayed elevated ABR thresholds at all tested frequencies compared to WT mice (**Fig. 7C**). These results demonstrate that ATL2 deficiency significantly increases auditory thresholds and impairs both the amplitude and latency of ABR waves, indicating a decline in hearing function.

## Discussion

Membrane shaping and remodeling of organelles is long thought to have critical physiological functions (Shibata et al., 2009). The significance of ATL initially drew attention because human mutations in ATL1 were linked to the pathogenesis of HSP (Zhao et al., 2001). Subsequent investigations revealed that ATL mediates membrane fusion of the ER by establishing the characteristic three-way junctions, generating an intricate membrane network (Hu et al., 2009)(Orso et al., 2009). As expected, defective ATL activities result in loss of complexity, and sometimes even fragmentation, of the ER in cells. Our results here explain how such defects cause specific retrograde degeneration of long axons in humans. In particular, compromised ATL action disturbs the homeostasis of ER export, and lipid synthesis pathways managed by SREBP appear to be most sensitive to the defects. Thus, the maintenance of ER dynamics by ATL serves to sense and surveil the health of the ER. Prolonged problems in ER network formation trigger a systemic freeze in lipid production.

As demonstrated previously (Niu et al., 2019), the impact of ATL deletion on ER export does not exhibit cargo specificity, both membrane and soluble cargoes are affected to different extents. In addition to SCAP trafficking, we noticed a possible decline in Sonic Hedgehog (Shh) signaling by Purkinje neurons, which can be explained by altered trafficking of both the ligand and the receptor. Shh secreted by Purkinje cells plays a key role in cerebellar development (Lewis et al., 2004). Shrinkage of the cerebellum, specifically slowed cell proliferation, was observed in ATL2 CKO of Purkinje neurons. Additional secreted clients that can be sensitive to ATL activity have yet to be identified in more physiological or pathological settings.

Notably, defects in lipid synthesis caused by ATL deletion in various cell types lead to divergent impacts in lipid distribution and membrane organization. In BG cells, key organelles, including the ER, mitochondria, and Golgi apparatus, exhibit a reduced volume upon loss of ATL. In contrast, the overall cell body size is not evidently altered. As a result, cell polarity and positioning are severely influenced. In Purkinje cells, which are known for their large size and complex cell surface (Busch and Hansel, 2023)(Yang et al., 2024), the plasma membrane area is significantly decreased when ATLs are deleted. These cells appear to be largely functional, but the strength in delivering developmental cues, particularly the Shh signals, is affected. In calyx-type synapses, the greatest shortage in membranes lies in synaptic vesicles. It is plausible that cells would distribute membrane resources to places where they are needed the most. Conversely, these places become more vulnerable when lipid synthesis is limited. How cells tune lipid distribution remains elusive.

Our KO analysis of ATL1 and ATL2 suggest that combined ATL activity is essential to cells. Neuroglia, such as BG cells, predominantly express ATL2, and its deletion causes devastating defects. Neurons like Purkinje cells and those composed of calyx-type synapses rely on both ATL1 and ATL2, as double KO results in an aggravated phenotype compared to single KO. These results provide a reasonable explanation for the discrepancy between humans and mice (i.e., different brain functions have different dependence on levels of ATL1 and ATL2). In humans, mutations of ATL1 alone are not severe enough to disturb brain development but compromise neuronal functions in cortical spinal motor neurons. These neurons likely demand normal activity of ATL1, and that of ATL2 can be suppressed by ATL1 via a dominant-negative effect. In mice, ATL1 mutations still show no significant defects in brain development, but mutations of both ATL1 and ATL2 can disrupt neuronal functions at the calyx of Held; the dependence of ATL1 is intrinsic to neurons and that of ATL2 is mostly in glia.

Patch-clamp recordings and EM analysis revealed similar defects at a mammalian central synapse, the calyx of Held, by deleting either ATL1 or ATL2. In both cases, synaptic vesicle docking at active zones was significantly diminished, resulting in a reduced RRP and impaired neurotransmitter release. Similarly, postsynaptic responses were attenuated. In addition, ATL deletion compromised both clathrin-dependent and clathrin-independent vesicle endocytosis. Direct shrinkage of the RRP offers a unique opportunity and a previously unavailable experimental model for understanding the plasticity of synaptic neurotransmission. Importantly, the observed reduction in docked synaptic vesicles directly impairs synaptic efficacy and timing precision, resulting in elevated auditory thresholds as analyzed by ABRs.

Functional studies of the calyx of Held using various ATL2 CKO mice confirmed that neurons rely heavily on glia to acquire lipid resources. Lipids are usually packed into lipoproteins, such as ApoE in the brain, secreted by glia, and then delivered to neurons (Yang et al., 2022). Therefore, a high-fat diet (HFD) may potentially alleviate some membrane shortage defects in neurons when ATL is defective. In fact, when pregnant mice were provided a HFD, cerebellar development defects could be partially rescued in SCAP CKO mice (Camargo et al., 2012). Impaired lipid metabolism in astrocytes, particularly levels of cholesterol, was found in cells derived from HSP patients (Mou et al., 2020). Similar strategies can be applied to neurodegenerative diseases other than HSP. Finally, because of the functional redundancy between ATL1 and ATL2, an ATL2 agonist, if available, may be beneficial to patients with defective ATL1 activity, a principle that has been proven in the case of mitochondrial outer membrane fusion, a similar membrane fusion process that is mediated by dynamin-like GTPase mitofusin (Guo et al., 2023).

## Method and materials

### Animals

ATL2 gene trap (*Atl2^wt/gt^*) mice were purchased from the International Mouse Phenotyping Consortium. ATL1 knockout and ATL1 K80A mutant mice were constructed by Cyagen using TALENs. Nes-Cre mice were a gift from Hong Zhang, Institute of Biophysics, Chinese Academy of Sciences. Cag-Flpo Atoh1-iCre and Cre-dependent reporter gene mice (*lsl-tdTomato*) were purchased from GemPharmatech. Pcp2-Cre and Gfap-Cre were purchased from Cyagen. Animals with conditional gene overexpression (*lsl-Atl1*, *lsl-Atl2*, *lsl-Cdh2*) were constructed by Cyagen using CRISPR-Cas9. All animal experiments were approved by the Institutional Committee at the Institute of Biophysics, Chinese Academy of Sciences

### Animal behavior test

Footprint test: the hindpaws of the mice were coated with non-toxic ink and guided to walk steadily on a white paper-lined runway (10 cm × 50 cm).

The rotarod test was performed to evaluate motor coordination and endurance in mice using an accelerating rotating rod (YLS-4C). During training, mice were placed on the rod at 4 rpm for 60 s, repeated three times with 10-min intervals. For formal testing, the rod accelerated linearly from 4 to 20 rpm over a 40 s period. Latency to fall (seconds) was automatically recorded, with a 180 s cutoff.

### Isolation and culture of mouse primary neuroglia

Neuroglia were isolated from the cerebrum cortex on postnatal day 4 (P4) according to the published protocol (Schildge et al., 2013). Briefly, P4 mouse pups were decapitated and the cortex dissected. Meninges were removed with fine forceps and the cortex hemispheres cut into small pieces. The cortex was incubated in 0.25% trypsin at 37°C for 30 min, centrifuged for 5 min at 300 x g, and the supernatant removed by decantation. The tissue was dissociated into a single cell suspension by adding 10 ml culture medium (DMEM, 10% FBS, p/s) and vigorously pipetted using a 10 ml plastic pipette. Cells were seeded on a culture dish (2 × 10^5^/cm^2^) coated with poly-D-lysine. The medium was changed every 2 days. Neuroglia were isolated from the cerebellum using the same method but cultured in DMEM/F12 supplemented with FGF-basic and B27.

### Cell lines and plasmids

COS-7 ATL2 and ATL3 double knockout cells were constructed previously in our lab (Niu et al., 2019). Cells were maintained in DMEM with 10% fetal bovine serum (Viva cell), 1% penicillin, and streptomycin. All cell lines were cultured under 5% CO_2_ at 37°C. SCAP-FLAG were gifts from Xiaowei Chen, Peking University.

### Transient siRNA-mediated knockdown

Small interfering RNA (siRNA) targeting ATL2 and ATL3 and non-specific siRNA control were purchased from RiboBio. The following siRNA sequences were used in this study: siATL2, 5′-GGAGCUAUCCUUAUGAACAUUCAUA-3′; siATL3, 5′-GGUUAGAGAUUGGAGUUUCCCUUAU-3′; Cells were transfected using Lipofectamine RNAiMAX (Invitrogen) according to the manufacturer’s protocol. All experiments were performed 48 hours after siRNA transfection. Knockdown efficiency was tested by western blot.

### Genotyping

Genomic DNA was extracted using lysis buffer (100 mM Tris-HCl pH 8.5, 6 mM EDTA, 0.2% SDS, 200 mM NaCl, 0.1 mg/ml proteinase K), precipitated using isopropanol, and dissolved in ddH_2_O. Genotyping was performed using Taq PCR mix (Mei5bio MF-002).

Primer pairs for genotyping were as follows:

gtKO-Forward (F2): TTCCTCAATCTCGCTCTCGC

gtKO-Reverse (R2): CCTGCGTGCAATCCATCTTG

fl-Forward (F1): GTAGTTATTCGGATCATCAGCTACAC

fl-Reverse (R1): GTAGTTATTCGGATCATCAGCTACAC

Cre-Forward: GTAGTTATTCGGATCATCAGCTACAC

Cre-Reverse: GTAGTTATTCGGATCATCAGCTACAC

atoh1-iCre-Forward: TGTCTGGTGTGGCTGATGAC

atoh1-iCre-Reverse: TTGGCACCATAGATCAGGCG

left arm-Forward (F3): CACTTGCTCTCCCAAAGTCGCTC

knockin-Reverse (R3): CCGTAAGTTATGTAACGCGGAACT

### RNA extraction and real-time PCR

After total RNA was isolated from cells using TRNzol (Thermo Fisher Scientific), cDNA was synthesized using a commercial kit (Genestar A230-10). RT-qPCR was performed using SYRB qPCR mix (Genestar ZA311) according to the manufacturer’s protocol. The target mRNA level was normalized to the GAPDH mRNA level. The following primers were used:

monkey-*FASN*-F: 5’-CAGGCGCTCAAGAAGGTGAT-3’

monkey-*FASN*-R: 5’-ATTGTACTCGGCGGAAGACG-3’

mouse-*CHKA*-F: 5’-TTGGCGATGAGCCTCGGAAAGT-3’

mouse-*CHKA*-R: 5’-GTGACCTCTCTGCAAGAATGGC-3’

mouse-*PYCT1B*-F: 5’-CTCTTCCACTCAGGTCACGCAA-3’

mouse-*PYCT1B*-R: 5’-CTCTCTGCCTCGTTCATCACAG-3’

mouse-*FASN*-F: 5’-CACAGTGCTCAAAGGACATGCC-3’

mouse-*FASN*-R: 5’-CACCAGGTGTAGTGCCTTCCTC-3’

mouse-*FDFT1*-F: 5’-GGATGTGACCTCCAAACAGGAC-3’

mouse-*FDFT1*-R: 5’-CAGACCCATTGAGTTGGCACAC-3’

mouse-*HMGCS1*-F: 5’-GGAAATGCCAGACCTACAGGTG-3’

mouse-*HMGCS1*-R: 5’-TACTCGGAGAGCATGTCAGGCT-3’

mouse-*SCD1*-F: 5’-GCAAGCTCTACACCTGCCTCTT-3’

mouse-*SCD1*-R: 5’-CGTGCCTTGTAAGTTCTGTGGC-3’

mouse-*SCD2*-F: 5’-GTCTGACCTGAAAGCCGAGAAG-3’

mouse-*SCD2*-R: 5’-GCAAGAAGGTGCTAACGCACAG-3’

*GAPDH*-F: 5’-CATCACTGCCACCCAGAAGACTG-3’

*GAPDH*-R: 5’-ATGCCAGTGAGCTTCCCGTTCAG-3’

### Western blotting

Proteins from tissues or cultured cells were lysed in cell lysis buffer (150 mM NaCl, 1% Triton X-100, 2 mM EDTA, and 50 mM Tris, pH 8.0) with proteinase inhibitor (Lableader) and analyzed by SDS-PAGE and western blot. Proteins were detected using the following specific antibodies: ATL1 (Sigma), ATL2 (Abcam, Proteintech), ATL3 (Proteintech) GFAP (Proteintech), NeuN (Sigma), beta-actin (Proteintech), BLBP (Abcam), beta-III-tubulin (Proteintech), calbindin (Abcam, Proteintech), FDFT1 (Abcam).

### Tissue section preparation

Fresh tissues were fixed in formalin at room temperature. For cryosectioning, the fixed tissues were dehydrated in 30% sucrose solution at 4°C. The tissue was embedded in OCT and the tissue block sliced into 10-μm slices, which were picked up with glass slides and stored in –80°C. For paraffin sections, the tissues were dehydrated with gradient alcohol and then embedded in wax. The wax tissue blocks were sliced 5-μm-thick. The slices were floated in water, picked up by the glass slides, and baked at 60°C before storing at room temperature.

### Immunofluorescence

Cell samples were fixed with 4% paraformaldehyde. For immunofluorescent staining, fixed cells were penetrated with 0.1% Triton X-100, blocked with 4% BSA, and incubated with primary antibody at room temperature for 1 hour. Samples were washed with PBS and incubated with fluorescence conjugated secondary antibody for 1 hour at room temperature. The samples were washed with PBS and mounted with DAPI-containing mounting medium. Mounted samples were stored at 4°C and observed under confocal microscopy within a week. For lipidtox staining, fixed cells were incubated with lipidtox dye for 15 min, washed with PBS, and immediately observed under confocal microscopy.

### Immunohistochemistry

Paraffin sections were baked at 70°C for 1 hour, and then put into xylene and graded alcohol for hydration. Antigen was repaired by heating the slices to 95°C for 15 min in antigen repairing buffer (Beyotime Biotechnology P0083). The slices were then incubated in 3% H_2_O_2_ to block endogenous peroxidase. The sections were blocked with 4% BSA, and then incubated with primary antibody and HRP-labeled secondary antibody. The tissue area was covered with DAB color developing solution, stained with hematoxylin, and rinsed with tap water. The slices were sealed with glycerol for microscopy.

### Cholesterol content assay

The cholesterol content of cells or tissues was analyzed using the cholesterol measurement kit according to the manufacturer’s protocol.

### Granule neuron migration

Images of cerebellum granule neuron migration were acquired as described previously (Benard et al., 2015). Briefly, cerebellums from mice at postnatal day 10 were sliced into acute cerebellar slices using Leica vibratomes and then stained with fluorescent dye (Invitrogen C2925 10 µM). Slices were transferred to the membrane of a Transwell insert and covered with culture medium. Images were taken every 15 min for up to 40 hours using a confocal microscope.

### Sample preparation for electronic microscopy

Tissue samples were fixed with 2.5% glutaraldehyde, and then immersed in 1% OsO_4_ and 1.5% potassium ferricyanide aqueous solution at 4°C for 1 h. After washing, they were incubated in 1% thiocarbohydrazide for 30 min, 1% OsO_4_ for 1 h, and 1% UA for 2 h. The tissues were dehydrated through graded alcohol into pure acetone. Samples were infiltrated in graded mixtures (3:1, 1:1, 1:3) of acetone and SPI-PON812 resin, then pure resin. Finally, tissues were embedded in pure resin.

### FIB-SEM and AutoCUTS-SEM

The serial block face was cut by a scanning electron microscope (FEI Helios Nanolab 600i dual-beam SEM) using a focused beam of gallium ions (30 kV, 0.43 nA) with a thickness of 20 nm, and imaged in immersion high magnification mode (CBS detector, 2 kV, 0.69 nA).

Automatic collector of ultrathin sections scanning electron microscopy (AutoCUTS-SEM) was performed as described previously for the 3D-ultrastructural study (Li et al., 2017). We collected sections with a thickness of 50 nm using an ultramicrotome (UC7, Leica, Germany) with the AutoCUTS (Zhenjiang Lehua Technology Co., Ltd, China) device. The serial sections were automatically acquired by a Helios Nanolab 600i dual-beam SEM (Thermo Fisher, USA) with automated imaging software (AutoSEE). The image parameters were: accelerating voltage of 2 kV, beam current of 0.69 nA, CBS detector.

### 3D reconstruction of cell organelles

After Z stacks were aligned by Ameria, organelles in EM images were marked using DeepContact (Liu et al., 2022) and cellSens software (Olympus). Marked organelle image stacks were rendered into 3D volumes and analyzed using Imaris v9.2 (Bitplane AG).

### Preparation of lipoprotein-deficient serum (LPDS)

Fetal bovine serum was adjusted to a density of 1.21 g/ml with KBr and centrifuged at 60,000 rpm for 27.6 h at 10°C in a Beckman type Ti70.1 rotor. Following centrifugation, the top fraction containing lipoproteins was removed and the bottom fraction containing LPDS was collected. The LPDS was dialyzed for 72 h at 4°C using 1 L PBS for every 10 ml of LPDS. The PBS was changed every 24 h. The LDPS was filtered into a sterile container using a 0.2-μm filter.

### Animals, slice preparation, and electrophysiology

Brain slices containing the medial nucleus of the trapezoid body (MNTB) region were prepared from mice of either sex on postnatal day 7-10 (p7-p10), and the transverse 200-μm-thick brainstem slices were prepared using a vibratome slicer (VT 1200s; Leica) as described previously (Wu et al., 2020). Brains were dissected and cut at 4°C in solution containing the following: 125 mM NaCl, 2.5 mM KCl, 0.05 mM CaCl_2_, 3 mM MgCl_2_, 25 mM glucose, 25 mM NaHCO_3_, 1.25 mM NaH_2_PO_4_, 0.4 mM L-ascorbic acid, 3 mM *myo*-inositol, and 2 mM Na-pyruvate (pH 7.4). Slices were incubated for 30 min at 37°C in solution containing the following: 125 mM NaCl, 2.5 mM KCl, 2 mM CaCl_2_, 1 mM MgCl_2_, 25 mM glucose, 25 mM NaHCO_3_, 1.25 mM NaH_2_PO_4_, 0.4 mM L-ascorbic acid, 3 mM *myo*-inositol, and 2 mM Na-pyruvate (pH 7.4).

For presynaptic recordings at the presynaptic terminal of the calyx-type synapse, the calcium current (ICa) and membrane capacitance (Cm) were measured in a whole-cell configuration using an EPC-10 amplifier (HEKA, Germany) with a software lock-in amplifier (1000 Hz sine wave, peak to peak voltage ≤ 60 mV). The presynaptic recording solution contained the following: 105 mM NaCl, 2.5 mM KCl, 2 mM CaCl_2_, 1 mM MgCl_2_, 25 mM glucose, 25 mM NaHCO_3_, 1.25 mM NaH_2_PO_4_, 0.4 mM L-ascorbic acid, 3 mM *myo*-inositol, 2 mM Na-pyruvate, 0.001 mM TTX, and 20 mM TEA-Cl (pH 7.4). The solution was maintained at 95% O_2_/5% CO_2_. The presynaptic pipette (3-5 MΩ) solution contained the following: 125 mM Cs-gluconate, 20 mM CsCl, 4 mM Mg-ATP, 10 mM Na_2_-phosphocreatine, 0.3 mM GTP, 10 mM HEPES, and 0.05 mM BAPTA (pH 7.4).

For postsynaptic recordings at the principal neuron of the calyx-type synapse, EPSCs were evoked by afferent fiber stimulation using a bipolar electrode positioned near the MNTB midline. Electrical pulses (0.1 Hz, <10 V) were delivered at an intensity 20% above the activation threshold. EPSCs were recorded with an EPC-10 amplifier using a pipette (2-3 MΩ) containing the following: 125 mM K-gluconate, 20 mM KCl, 10 mM Na_2-_phosphocreatine, 4 mM Mg-ATP, 0.3 mM GTP, 10 mM HEPES, and 0.5 mM EGTA (pH 7.2 adjusted with KOH).

Series resistance (<10 MΩ) was compensated by 95% with a 10 µs lag. A paired stimulus (20 ms inter-stimulus interval) were applied for paired-pulse experiments to elicit paired EPSCs. The paired-pulse ratio (PPR) was calculated as the amplitude ratio of the second EPSC to the first EPSC (EPSC_2_/EPSC_1_).

### Data collection and statistical analyses

The initial endocytosis rate (Rate_endo_) was measured within 2 s after depolarization. The percentage of residual capacitance 15 s (ΔCm_15s_%) or 30 s (ΔCm_30s_%) after depolarization was measured to represent capacitance recovery.

For recordings of ICa, capacitance changes, and postsynaptic responses at calyx-type synapses, data were collected from 5-13 calyceal terminals obtained from 3-7 mice. Data were analyzed using PatchMaster (HEKA), Igor 9 (WaveMetrics), GraphPad Prism 9, and ZEN (Zeiss).

### Tissue lipid extraction and lipidomics analyses

Lipids were extracted from frozen tissues using a modified version of the Bligh and Dyer’s method as described previously (Lam et al., 2022). Briefly, tissues were homogenized in 750 µl of chloroform:methanol:MilliQ H_2_O (3:6:1 v/v/v). The homogenate was then incubated for 1 h at 4°C at 1500 rpm. At the end of the incubation, 350 µl of deionized water and 250 µl of chloroform were added to induce phase separation. The samples were then centrifuged and the lower organic phase containing lipids was extracted into a clean tube. Lipid extraction was repeated once by adding 450 µl of chloroform to the remaining aqueous phase, and the lipid extracts were pooled into a single tube and dried in the SpeedVac under OH mode. Samples were stored at –80°C until further analysis.

Lipidomic analyses were conducted at LipidALL Technologies using a ExionLC-AD coupled with Sciex QTRAP 6500 PLUS as reported previously (Lam et al., 2021). Separation of individual lipid classes of polar lipids by normal phase (NP)-HPLC was achieved using a TUP-HB silica column (i.d. 150×2.1 mm, 3 µm) under the following conditions: mobile phase A (chloroform:methanol:ammonium hydroxide, 89.5:10:0.5) and mobile phase B (chloroform:methanol:ammonium hydroxide:water, 55:39:0.5:5.5). MRM transitions were set-up for comparative analysis of various polar lipids. Individual lipid species were quantified by reference to spiked internal standards. We obtained d9-PC32:0(16:0/16:0), d9-PC36:1p(18:0p/18:1), d7-PE33:1(15:0/18:1), d9-PE36:1p(18:0p/18:1), d31-PS(d31-16:0/18:1), d7-PA33:1(15:0/18:1), d7-PG33:1(15:0/18:1), d7-PI33:1(15:0/18:1), C17-SL, d5-CL72:8(18:2)4, Cer d18:1/15:0-d7, d9-SM d18:1/18:1, C8-GluCer, C8-GalCer, d3-LacCer d18:1/16:0, Gb3 d18:1/17:0, d7-LPC18:1, d7-LPE18:1, C17-LPI, C17-LPA, C17-LPS, C17-LPG, d17:1 Sph, and d17:1 S1P from Avanti Polar Lipids. GM3-d18:1/18:0-d3 was purchased from Matreya LLC. Free fatty acids were quantitated using d31-16:0 (Sigma-Aldrich) and d8-20:4 (Cayman Chemicals).

Glycerol lipids including diacylglycerol (DAG) and triacylglycerol (TAG) were quantified using a modified version of reverse phase HPLC/MRM (Shui et al., 2010). Separation of neutral lipids was achieved on a Phenomenex Kinetex-C18 column (i.d. 4.6×100 mm, 2.6 µm) using an isocratic mobile phase containing 100:100:4 chloroform:methanol:0.1 M ammonium acetate (v/v/v) at a flow rate of 300 µl for 10 min. Levels of short-, medium-, and long-chain TAGs were calculated by reference to spiked internal standards of TAG(14:0)3-d5,TAG(16:0)3-d5 and TAG(18:0)3-d5 obtained from CDN isotopes, respectively. DAGs were quantified using d5-DAG17:0/17:0 and d5-DAG18:1/18:1 as internal standards (Avanti Polar Lipids).

Free cholesterols and cholesteryl esters were analyzed under atmospheric pressure chemical ionization (APCI) mode on a Jasper HPLC coupled to Sciex 4500 MD as described previously using d6-cholesterol and d6-C18:0 cholesteryl ester (CE) (CDN isotopes) as internal standards (Shui et al., 2011).

### Cell lipid extraction and lipidomics analyses

The cell pellets were mixed with beads and 200 μl H_2_O and underwent three freeze–thaw cycles. The samples were mixed with 480 μl of extraction solution (MTBE:MeOH, 5:1 (v/v)) containing deuterated internal standards. The samples were incubated for 1 h at –40°C to precipitate proteins and then centrifuged at 900×g for 15 min at 4°C. The supernatant was removed and evaporated to dryness. Dry extracts were reconstituted in 100 μl of DCM:MeOH (1:1, v/v) and then centrifuged at 13,800×g. A total of 75 μl of supernatant was collected for analysis.

For lipids, LC-MS/MS analyses were performed using an UHPLC system (Vanquish, Thermo Fisher Scientific) with the Phenomenex Kinetex C18 (2.1 mm × 100 mm, 2.6 μm) coupled to an Orbitrap Exploris 120 mass spectrometer (Orbitrap MS, Thermo). Mobile phase A was H2O/ACN (6:4, v/v) with 10 mM HCOONH4, and mobile phase B was IPA/ACN (9:1, v/v) with 10 mM HCOONH_4_. The injection volume was 2 μl. The Orbitrap Exploris 120 mass spectrometer was used for its ability to acquire MS/MS spectra under the control of the acquisition software (Xcalibur, Thermo). In this mode, the acquisition software continuously evaluates the full scan MS spectrum. The ESI source conditions were set as follows: sheath gas flow rate 30 Arb; Aux gas flow rate 10 Arb; capillary temperature 320°C; full MS resolution 60,000; MS/MS resolution 15,000; collision energy, SNCE 15/30/45; spray voltage 3.8 kV (positive) or –3.4 kV (negative).

### Quantification and statistical analysis

Statistical analyses were performed using Prism 9.0 (GraphPad). All experiments were performed three times. Data are provided as mean ± SEM or mean ± SD. The data analysis method was one-way ANOVA with Dunnett’s *post hoc* test or unpaired Student’s t test.

## Acknowledgements

We thank Dr. Haitao Wu for Cre mice and helpful discussion, Dr. Alicia Prater for proofreading, and Xixia Li, Zhongshuang Lv, and Can Peng, Liqing Liu, Yun Feng, and Chunliu Liu at the Center for Biological Imaging (CBI), Institute of Biophysics, Chinese Academy of Science for technical support. JH is supported by grants from the National Natural Science Foundation of China (32230024 and 92254305) and Project for Young Scientists in Basic Research (YSBR-075) of the Chinese Academy of Sciences. LX is supported by a grant from the Science and Technology Innovation 2030 – Brain Science and Brain-Inspired Intelligence Project (2021ZD0201301), the National Key Research & Development Program of China (2022YFC3602700&2022YFC3602702), and the National Natural Science Foundation of China (32170688).

---

**Figure S1.**
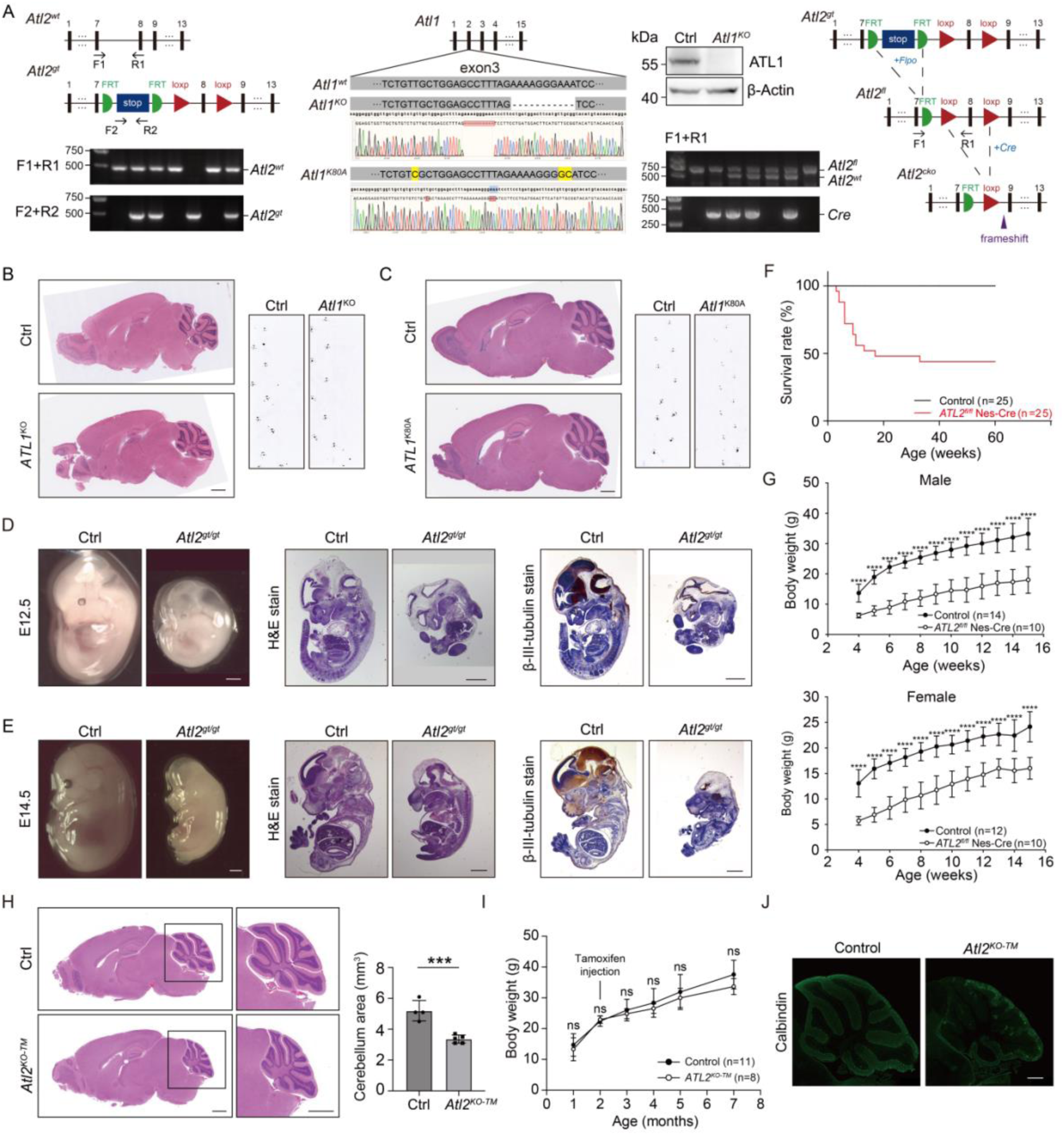
ATL2 is essential for cerebellar development. **A:** Left, schematic of the ATL2 mutant allele (*ATL2^gt^*) and PCR genotyping. Black blocks represent exons with specific numbers indicated. F1, R1, F2, and R2 represent genotyping primers. Middle, schematic of the ATL1 knockout, ATL1 K80A mutant mouse, their sequencing result and western blot analysis of ATL1 expression in control and *ATL1^KO^* mouse brain. Right, schematic of construction of the conditional knockout mouse and PCR genotyping. *ATL2^gt^* mice were crossed with the Flpo mouse to remove the stop sequence and produce *ATL2^fl^* mice, then *ATL2^fl^*mice were crossed with a Cre mouse to generate conditional knockout animals. Detailed information on the *ATL2^gt^* mouse: https://www.mousephenotype.org/data/alleles/MGI:1929492/tm1a(EUCOMM)Hmgu?alleleSymbol=tm1a(EUCOMM)Hmgu#targetingVector. **B-C:** H&E staining (left) and footprint test (right) of *ATL1^KO^* (B) and *ATL1^K80A^* (C) mice and their controls (scale bars, 1 mm). **D-E:** Images (left), H&E staining (middle), and β-III-tubulin IHC staining (right) of E12.5 (D) and E14.5 (E) mice (scale bar, 1 mm). **F-G:** Survival rate (F) and body weight (G) of control and *ATL2^fl/fl^* Nes-Cre mice. **H:** H&E staining (left) and area (right) of the cerebellum of control and *ATL2^KO-TM^* mice (scale bar, 1 mm). **I:** Body weight of control and *ATL2^KO-TM^* mice. **J:** Calbindin staining of control and *ATL2^KO-TM^*mice (scale bar, 0.5 mm). Data are represented as mean ± SD. The statistical significance of mean values was analyzed using an unpaired Student’s t test, \*\*\*\**p* < 0.0001, \*\*\**p* < 0.001, ns, not significant.

**Figure S2.**
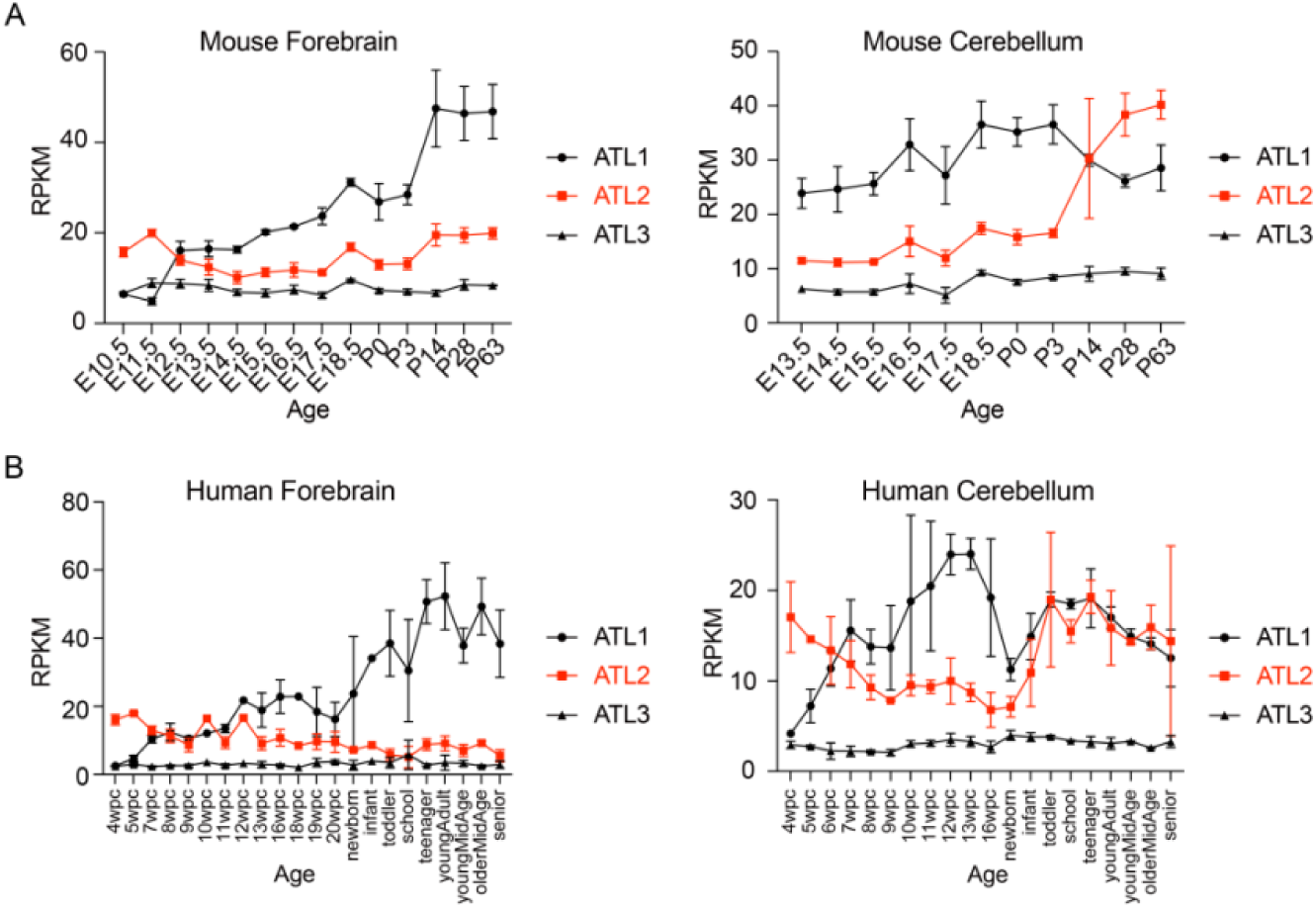
Levels of ATL transcripts in brain tissues. **A-B:** RNA level of three ATLs in mouse (A) and human (B) forebrain and cerebellum. Data are represented as mean ± SD.

**Figure S3.**
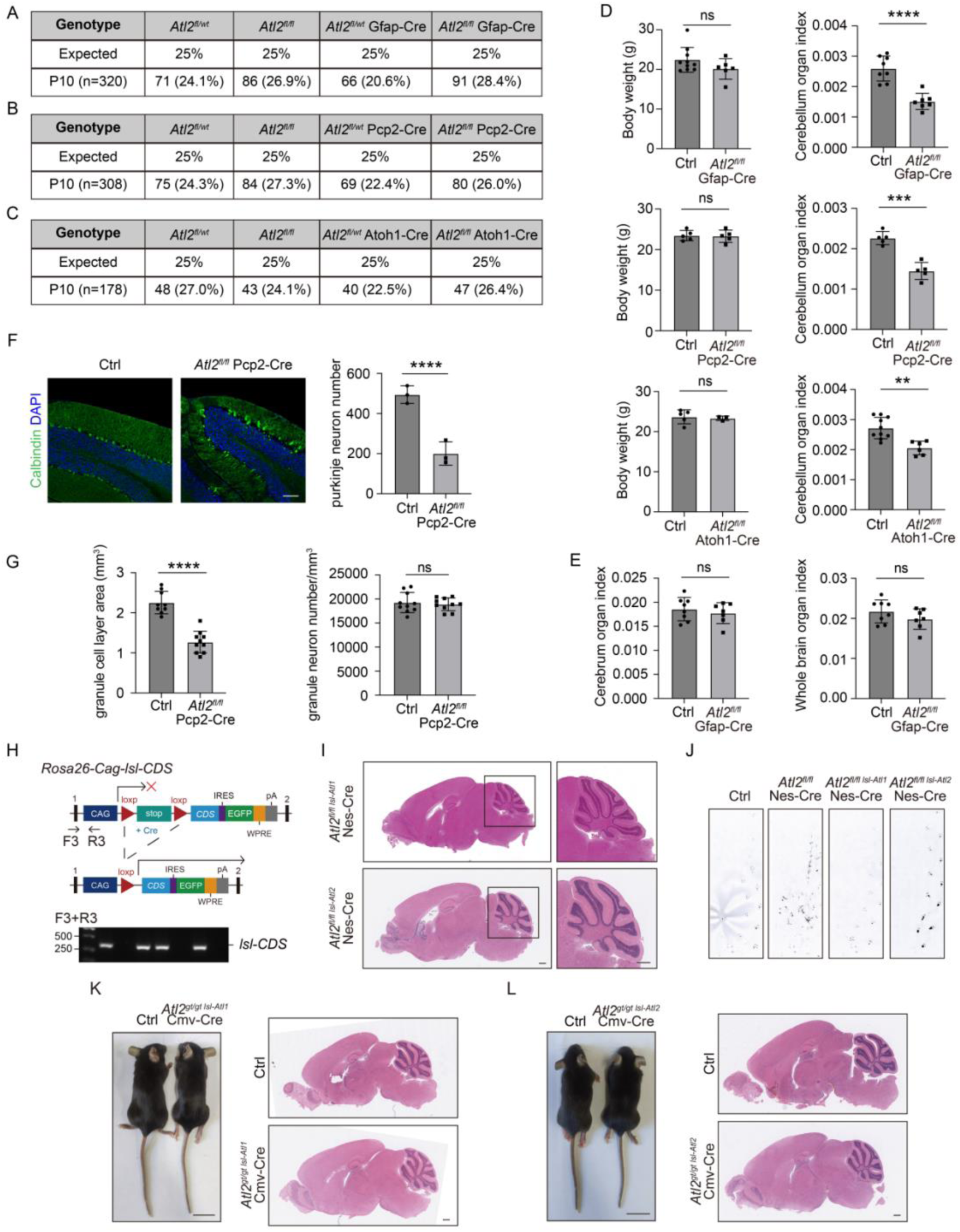
Loss of ATL2 affects different cell types in the cerebellum. **A-C:** Distributions of offspring from the intercross between (A) *ATL2^fl/fl^* and *ATL2^fl/wt^*Gfap-Cre mice, (B) *ATL2^fl/fl^* and *ATL2^fl/wt^* Pcp2-Cre mice, (C) *ATL2^fl/fl^* and *ATL2^fl/wt^* Atoh1-Cre mice. **D:** Body weight and cerebellum organ index of *ATL2^fl/fl^* Gfap-Cre, *ATL2^fl/fl^* Pcp2-Cre, *ATL2^fl/fl^* Atoh1-Cre mice, and their controls. **E:** Cerebellum and whole brain organ index of control and *ATL2^fl/fl^* Gfap-Cre mice. **F:** Calbindin immunofluorescent staining and cell count of cerebellum Purkinje neurons (scale bar, 100 μm). **G:** Area and cell density of the cerebellum granule cell layer of control and *ATL2^fl/fl^* Pcp2-Cre mice. **H:** Schematic of gene conditional overexpression mice. Conditional gene expression elements were inserted between exons 1 and 2 of the Rosa26 safe harbor locus. Lower panel, PCR genotyping. **I:** H&E staining of sagittal sections of cerebellum from *ATL2^fl/fl^* Nes-Cre mice overexpressing ATL1 or ATL2 (scale bar, 0.5 mm). **J:** Footprint test of control, *ATL2^fl/fl^* Nes-Cre, and *ATL2^fl/fl^* Nes-Cre mice overexpressing ATL1 or ATL2. **K-L:** Image (K) and H&E staining of sagittal cerebellum sections (L) of *ATL2^gt/gt^*mice overexpressing ATL1 or ATL2 (scale bar: 2 cm in image, 0.5 mm in H&E staining). Data are represented as mean ± SD. The statistical significance of mean values was analyzed using an unpaired Student’s t test, \*\*\*\**p* < 0.0001, \*\*\**p* < 0.001, \*\**p* < 0.01, ns, not significant.

**Figure S4.**
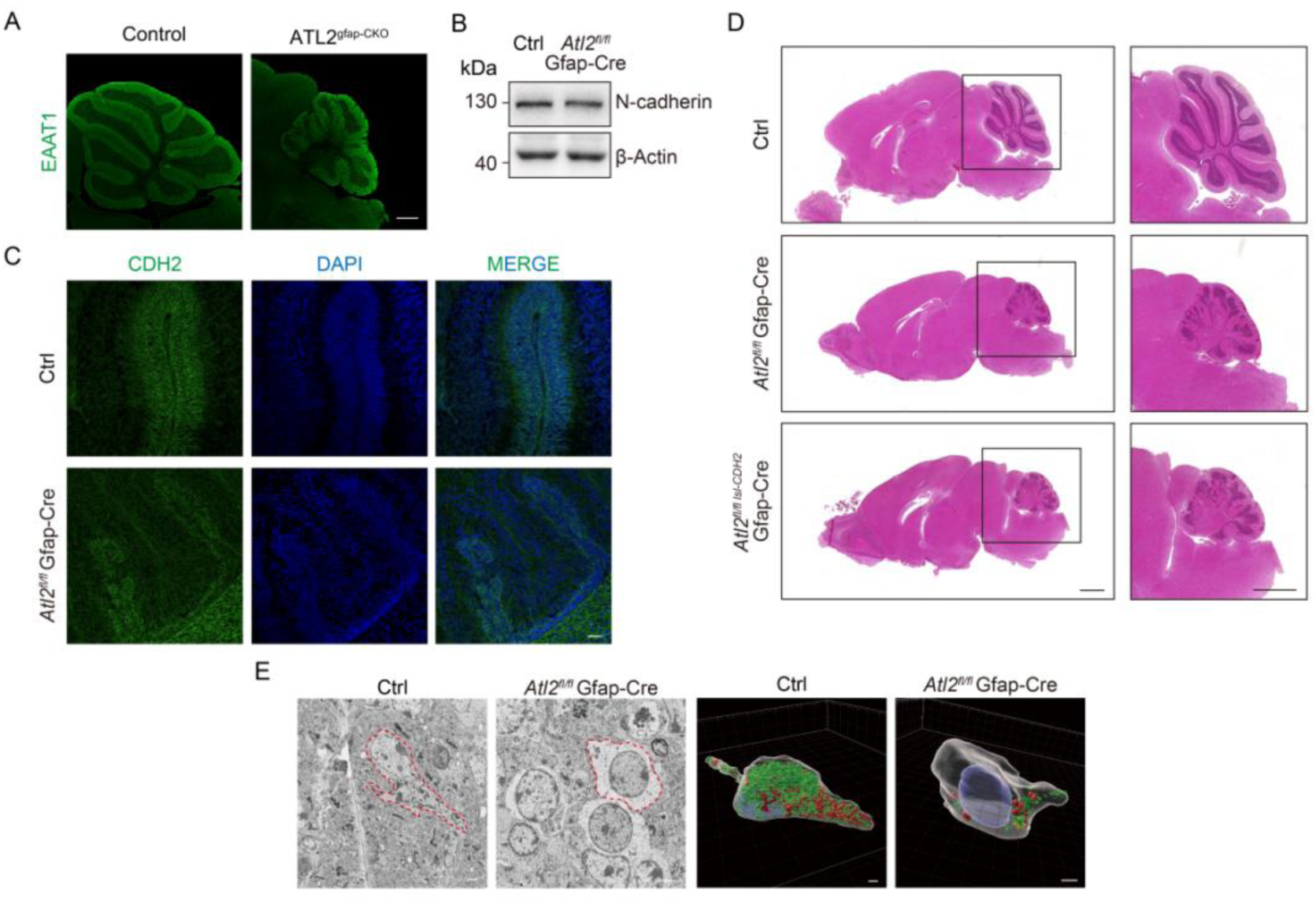
Effect of ATL2 knockout on cerebellar neuroglia. **A:** EAAT1 immunofluorescent staining of control and *ATL2^fl/fl^*Gfap-Cre mouse cerebellum (scale bar: 0.5 mm). **B:** Western blot analysis of CDH2 expression in control and *ATL2^fl/fl^* Gfap-Cre mouse cerebellum. **C:** CDH2 immunofluorescent staining of control and *ATL2^fl/fl^*Gfap-Cre mouse cerebellum (scale bar: 50 μm). **D:** H&E staining of brains of control, *ATL2^fl/fl^* Gfap-Cre, and *ATL2^fl/fl^*Gfap-Cre mice overexpressing CDH2 (scale bar: 1 mm). **E:** Electron microscopy image and 3D reconstruction of Bergmann glia from serial sections of mouse cerebellum (scale bar: 2 μm).

**Figure S5.**
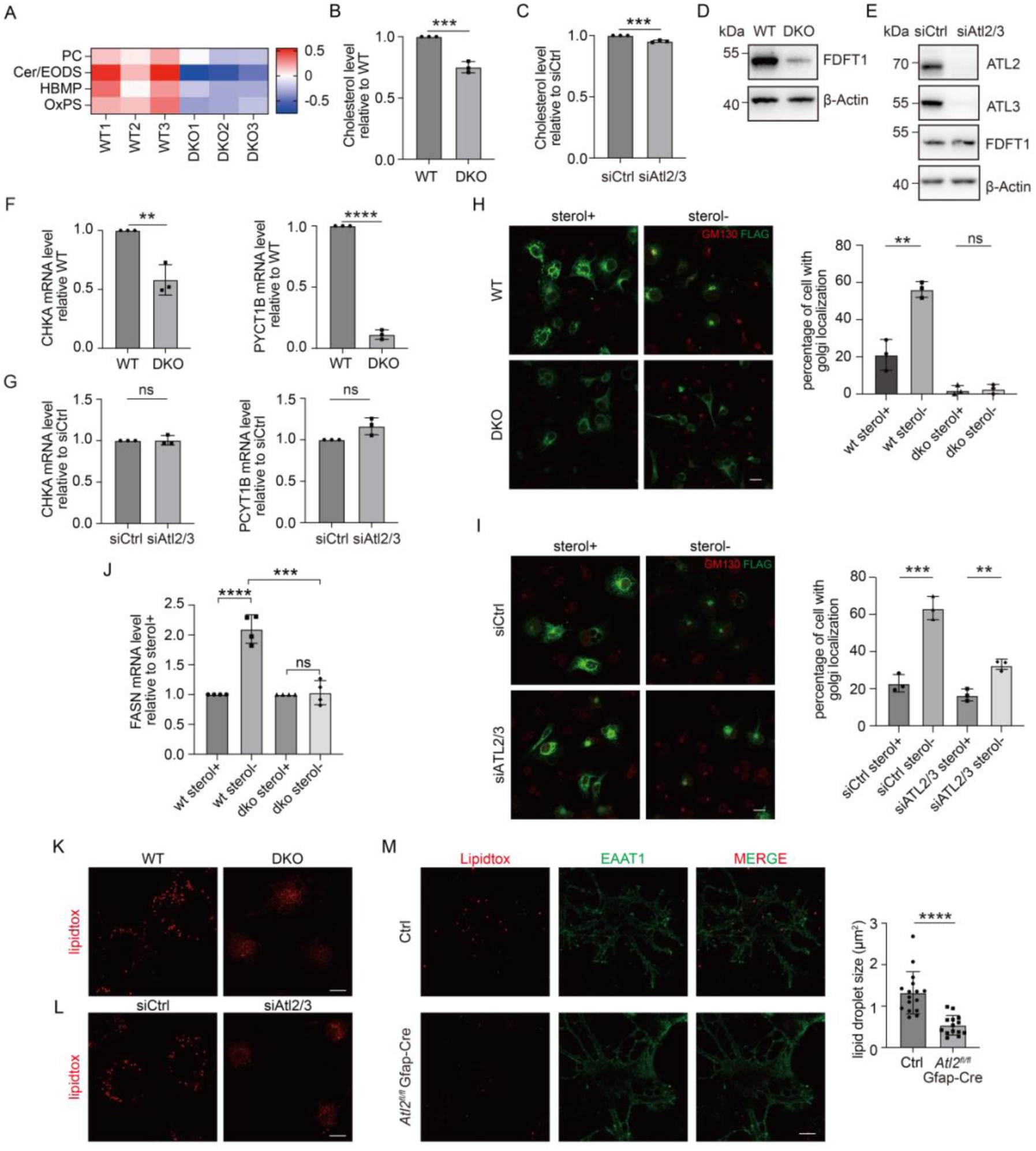
Knockout of ATL2 disrupts lipid homeostasis. **A:** Lipidomics analysis of WT and ATL2/3 DKO COS7 cells. **B-C:** Relative change in the cholesterol level in ATL2/3 DKO (B) and ATL2/3 knockdown (C) COS7 cells. **D-E:** Western blot analysis of FDFT1 in ATL2/3 DKO (D) and ATL2/3 knockdown (E) COS7 cells. **F-G:** RT-PCR analysis of CHKA and PCYT1B mRNA levels in ATL2/3 DKO (F) and ATL2/3 knockdown (G) COS7 cells. **H-I:** ATL2/3 DKO (H) and ATL2/3 knockdown (I) COS7 cells transfected with SCAP-Flag were cultured in sterol-deprived conditions or treated with1 μg/ml 25-HC. Cells were fixed and subjected to confocal microscopy following staining with antibodies against GM130 (red) and Flag (green) (scale bar: 20 μm). **J:** RT-PCR analysis of FASN mRNA levels in WT and ATL2/3 DKO COS7 cells cultured in sterol-deprived conditions or treated with1 μg/ml 25-HC. **K-L:** Lipidtox staining of ATL2/3 DKO (K) and ATL2/3 knockdown (L) COS7 cells (scale bar: 10 μm). **M:** Lipidtox staining of neuroglia isolated from the cerebellum of control and *ATL2^fl/fl^* Gfap-Cre mice. EAAT1 was stained to indicate neuroglial cells (scale bar: 10 μm). Data are represented as mean ± SD. The statistical significance of mean values was analyzed using an unpaired Student’s t test, \*\*\*\**p* < 0.0001, \*\*\**p* < 0.001, \*\**p* < 0.01, ns, not significant.

**Figure S6.**
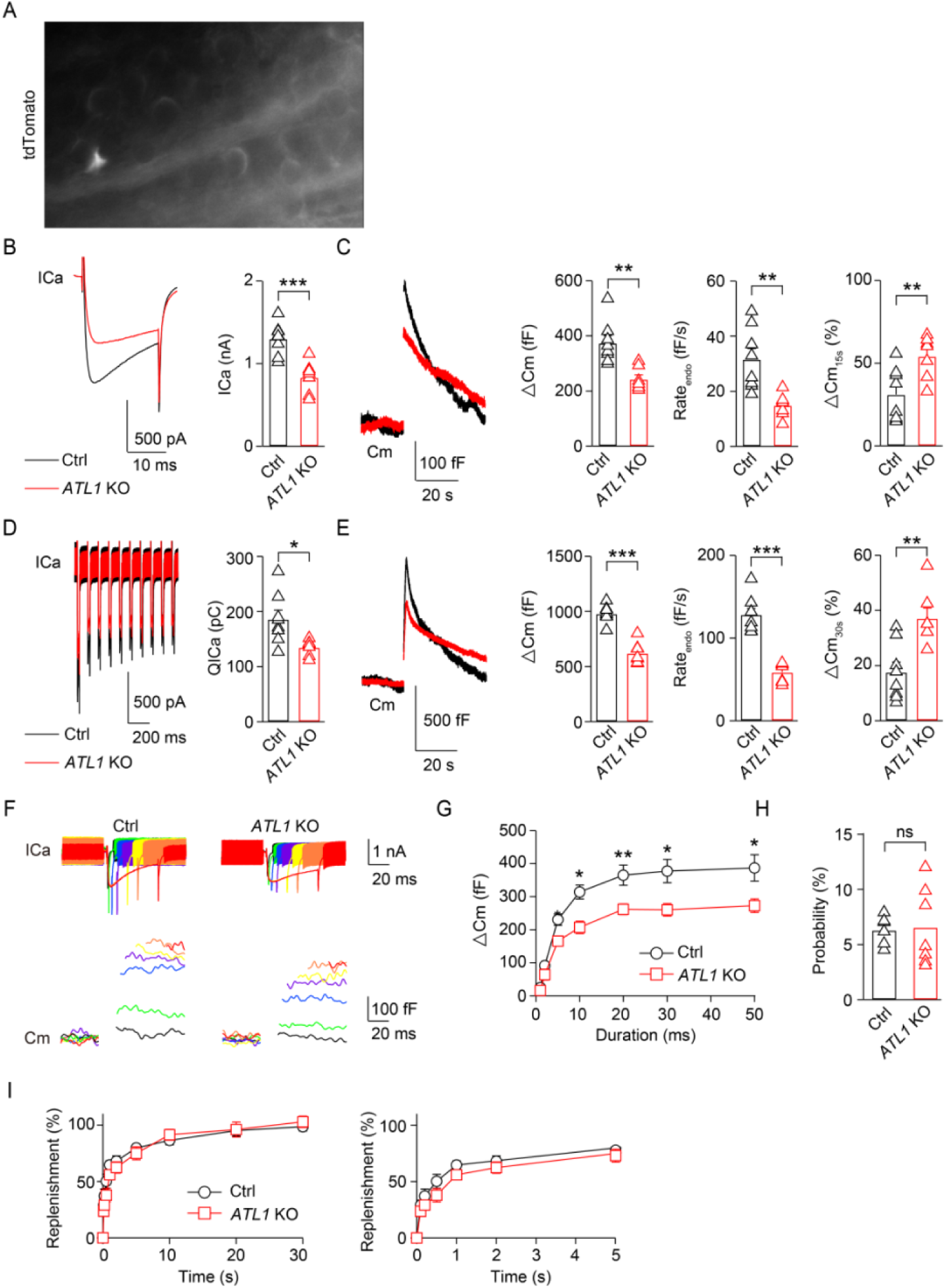
Deletion of ATL1 inhibits presynaptic ICa, exo-endocytosis, and RRP size. **A**: Positive *Tdtomato* reporter expression indicated that Atoh1-Cre was expressed in the MNTB region. **B**: Left, averaged presynaptic ICa induced by depol_20ms_ in control (black) and ATL1 KO (red) mice. Right, statistics for presynaptic ICa induced by depol_20ms_ in control (black, n = 8) and ATL1 KO (red, n = 7) mice. **C**: Left, averaged Cm induced by depol_20ms_ in control (black) and ATL1 KO (red) mice. Right, statistics for the ΔCm (C), Rate_endo_ (D), and ΔCm_15s_ in control (black, n = 8) and ATL1 KO (red, n = 7) mice. **D**: Left, averaged presynaptic ICa induced by depol_20msx10_ (arrow) in control (black) and ATL1 KO (red) mice. Right, statistics for presynaptic QICa induced by depol_20msx10_ in control (black, n = 8) and ATL1 KO (red, n = 7) mice. **E**: Left, averaged Cm induced by depol_20msx10_ in control (black) and ATL1 KO (red) mice. Right, statistics for the ΔCm (C), Rate_endo_ (D), and ΔCm_30s_ in control (black, n = 8) and ATL1 KO (red, n = 7) mice. **F**: Sampled ICa (top) and Cm (bottom) induced by 1 ms (black), 2 ms (green), 5 ms (blue), 10 ms (purple), 20 ms (yellow), 30 ms (orange), and 50 ms (red) depolarization pluses from –80 to +10 mV in control and ATL1 KO mice. **G**: Relationship between ΔCm and the duration of depolarization pulses in control (n = 6 for each data point; black) and ATL1 KO (n = 7 for each data point; red) mice. **H**: Statistics for the release probability measured by the percentage of RRP release induced by 1-ms depolarization pulses from –80 to +10 mV in control (black) and ATL1 KO (red) mice using an unpaired Student’s t test; ns, not significant. **I**: Cm induced by a 20-ms depolarization applied at various intervals after the conditional stimulus (depol_20ms_) in control (black) and ATL1 KO (red) mice. Left and right panels show the same data at different scales. Data are represented as mean ± SEM. The statistical significance of mean values was analyzed using an unpaired Student’s t test, \*\*\**p* < 0.001, \*\**p* < 0.01, \**p* < 0.05, ns, not significant.

